# OCA-B promotes pathogenic maturation of stem-like CD4^+^ T cells and autoimmune demyelination

**DOI:** 10.1101/2023.11.29.569210

**Authors:** Erik P. Hughes, Amber R. Syage, Elnaz Mirzaei Mehrabad, Thomas E. Lane, Benjamin T. Spike, Dean Tantin

## Abstract

Stem-like T cells selectively contribute to autoimmunity, but the activities that promote their pathogenicity are incompletely understood. Here, we identify the transcription coregulator OCA-B as a driver of the pathogenic maturation of stem-like CD4^+^ T cell to promote autoimmune demyelination. Using two human multiple sclerosis (MS) datasets, we show that *POU2AF1*, the gene encoding OCA-B, is elevated in CD4^+^ T cells from MS patients. We show that T cell-intrinsic OCA-B loss protects mice from experimental autoimmune encephalomyelitis (EAE) while preserving responses to viral CNS infection. In EAE models driven by antigen reencounter, OCA-B deletion nearly eliminates CNS infiltration, proinflammatory cytokine production and clinical disease. OCA-B-expressing CD4^+^ T cells of mice primed with autoantigen express an encephalitogenic gene program and preferentially confer disease. In a relapsing-remitting EAE model, OCA-B loss protects mice specifically at relapse. During remission, OCA-B promotes the expression of *Tcf7*, *Slamf6*, and *Sell* in proliferating CNS T cell populations. At relapse timepoints, OCA-B loss results in both the accumulation of an immunomodulatory CD4^+^ T cell population expressing *Ccr9* and *Bach2*, and loss of pro-inflammatory gene expression from Th17 cells. These results identify OCA-B as a driver of pathogenic CD4^+^ T cells.

## Introduction

2.8 million individuals worldwide suffer from multiple sclerosis (MS), with almost 1 million cases in the United States (1). Despite recent advances, the development of new therapies that block MS while sparing mechanisms that prevent infection and viral recrudescence remains a critical goal. Genome-wide association studies (GWAS) identify the *HLA* loci as the most important determinant of MS pathogenesis, emphasizing the impact of T cells (2, 3). The association with the major histocompatibility complex (MHC) class II locus in particular indicates that myelin-specific CD4^+^ T cells are key regulators of disease.

The predominant preclinical model of MS is experimental autoimmune encephalomyelitis (EAE), in which laboratory animals are immunized with myelin proteins or peptides to activate antigen-specific CD4^+^ T cells that infiltrate the central nervous system (CNS) and mediate disease. EAE within the C57BL/6 mouse background establishes chronic disease in which Th1 and Th17 effector CD4^+^ T cell subsets are pathogenetic mediators (4, 5). After accessing the CNS, these cells secrete interferon-gamma (IFNγ), interleukin-17 (IL-17), granulocyte macrophage stimulating-factor (GM-CSF) and other inflammatory mediators and chemoattractants that recruit the cell types responsible for driving demyelination (6–8).

Myelin-reactive T cells from the blood of MS patients show stem/memory-like properties, with reduced need for co-stimulation (9–12). Consistent with these findings, autoreactive and bystander memory-phenotype CD4^+^ T cells preferentially transfer demyelinating disease in mouse models (13–15). While these studies have identified vital CD4^+^ T cell populations in EAE, further research on the mechanisms sustaining these populations and driving their pathogenic activity is required to develop more precise therapies.

Oct1, a POU domain transcription factor encoded by *Pou2f1*, has been linked to autoimmune disease. GWAS studies have identified Oct1 binding site polymorphisms that associate with a predisposition for autoimmune diseases, including MS (16–19). In CD4^+^ T cells, Oct1 promotes the development of both CD4^+^ memory (20) and EAE pathogenesis and demyelination (21). In contrast, Oct1 loss preserves immune responses to the neurotrophic viral infection (21). These observations highlight a promising pathway which when targeted may limit autoimmune CNS demyelination without the broad immune suppression elicited by many current therapeutics. However, Oct1 is widely expressed and regulates both embryonic development and somatic stem cell function (22–24), making it a poor therapeutic target.

The lymphoid-restricted transcription cofactor OCA-B regulates transcription by docking with Oct1, and in B cells an additional related factor, Oct2 (25). In B cells, OCA-B is essential for B cell maturation and germinal center formation (26–28). In contrast to Oct1, OCA-B whole-body knockouts are viable and fertile (28). Polymorphisms in the *OCAB* (*POU2AF1*) locus are associated with autoimmune diseases, including MS (29, 30). Although its expression is 50-100 fold lower in T cells compared to B cells (31), targeting either OCA-B by genetic deletion in T cells or its downstream functions with a membrane-permeable peptide mimic suppresses type-1 diabetes (32). OCA-B interacts with target genes such as *Il2, Ifng, Il21, Csf2 (Gmcsf), Tnfrsf4 (Ox40), Icos* and *Ctla4*, but is dispensable for their expression following primary activation of T cells. Instead, OCA-B regulates these targets under a narrow range of conditions, for example following secondary stimulation in culture (20). Many of these targets are directly implicated in MS pathogenesis (6–8, 33–39). OCA-B controls expression of *Tbx21* (*Tbet*), which encodes a key regulator of Th1 effector responses, but also promotes Th17 differentiation and may interact with RORγt, a master regulator of Th17 cells (20, 40, 41). In the context of MS, the only prior OCA-B work used whole body knockouts and showed partial protection in chronic C57BL/6 EAE models (41). Cumulatively, these findings indicate that OCA-B plays poorly defined but possibly significant roles in MS pathogenesis.

Here, we show that OCA-B within the T cell compartment drives EAE by promoting pathogenic maturation of stem-like CD4^+^ T cells. In C57BL/6 mice, T cell OCA-B loss significantly protects mice from chronic EAE while preserving responses to the neurotrophic coronavirus JHMV. Using passive transfer assays in which reactive T cells are primed with antigen in donor mice, polarized in culture, then reencounter antigen in naïve recipients, we show that OCA-B is critical for the infiltration of pathogenic cells into the CNS and the manifestation of clinical disease. Using an OCA-B reporter mouse, we find that OCA-B expressing CD4^+^ T cells taken from MOG peptide primed mice preferentially display pathogenic stem-like gene expression patterns and preferentially transfer demyelinating disease, highlighting OCA-B as both a marker and promoter of encephalitogenic autoimmune CD4^+^ T cell activity. Intriguingly, T cell conditional OCA-B knockout in EAE models on the autoimmune-prone non-obese diabetic (NOD) background, which develop relapsing-remitting disease (42), specifically protects against relapse. Single-cell transcriptomic analysis of CD3͛^+^, CNS-infiltrating T cells at remission and relapse indicates that OCA-B promotes disease relapse through control of stem-like T cell populations, driving them to pathogenic Th17 differentiation. Cumulatively, these findings mark OCA-B as a critical mediator of the pathogenic maturation of autoreactive T cells, and a promising potential therapeutic target.

## Results

### Elevated expression of OCAB in CD4+ T cells from secondary-progressive human MS lesions

To determine if the gene encoding OCA-B is elevated in human MS samples, we mined high-throughput single-nucleus RNA-seq data of glial and immune cells from a recent study using single-nucleus RNA-seq to evaluate gene expression in brain tissue from 54 MS patients and 26 non-MS controls (43). The N=1 RRMS sample showed far higher *POU2AF1* expression compared to CTR (Figure 1A). There were more primary-progressive (PPMS) samples, and interestingly only one sample showed significant expression, perhaps reflecting imperfections in diagnoses. Most dramatically, a subset of secondary-progressive (SPMS) samples showed significantly elevated OCA-B expression (Figure 1A). We then reclustered CD4^+^ cell nuclei from this dataset. Uniform Manifold Approximation and Projection (UMAP) feature plots revealed that the elevated *POU2AF1* found in T cell nuclei from SPMS patients was confined to one specific CD4^+^ cluster that also expresses the pathogenic marker *IL1R1* (Figure 1B). This cluster was also marked by the expression of the T cell activation marker *CD44* and the transcription factors *LEF1*, *ETS1*, *BCL6*, and *MAF* (Supplemental Figure 1A). To assess if elevated *POU2AF1* expression was restricted to CNS infiltrating T cells, we analyzed recent bulk RNAseq dataset (44) comparing peripheral blood memory T effector cells (“Teff”, CD4^+^CD25^low/neg^CD127^hi^CD45RO^+^) from 20 RRMS patients and 20 healthy controls (“HC”). Similar to CNS T cells, *POU2AF1* expression in peripheral CD4^+^ T cells was found elevated in MS patients compared to non-MS controls (Supplemental Figure 1B). These findings identify elevated *OCAB* mRNA expression in T cells from MS patients across multiple datasets.

**Figure 1.**
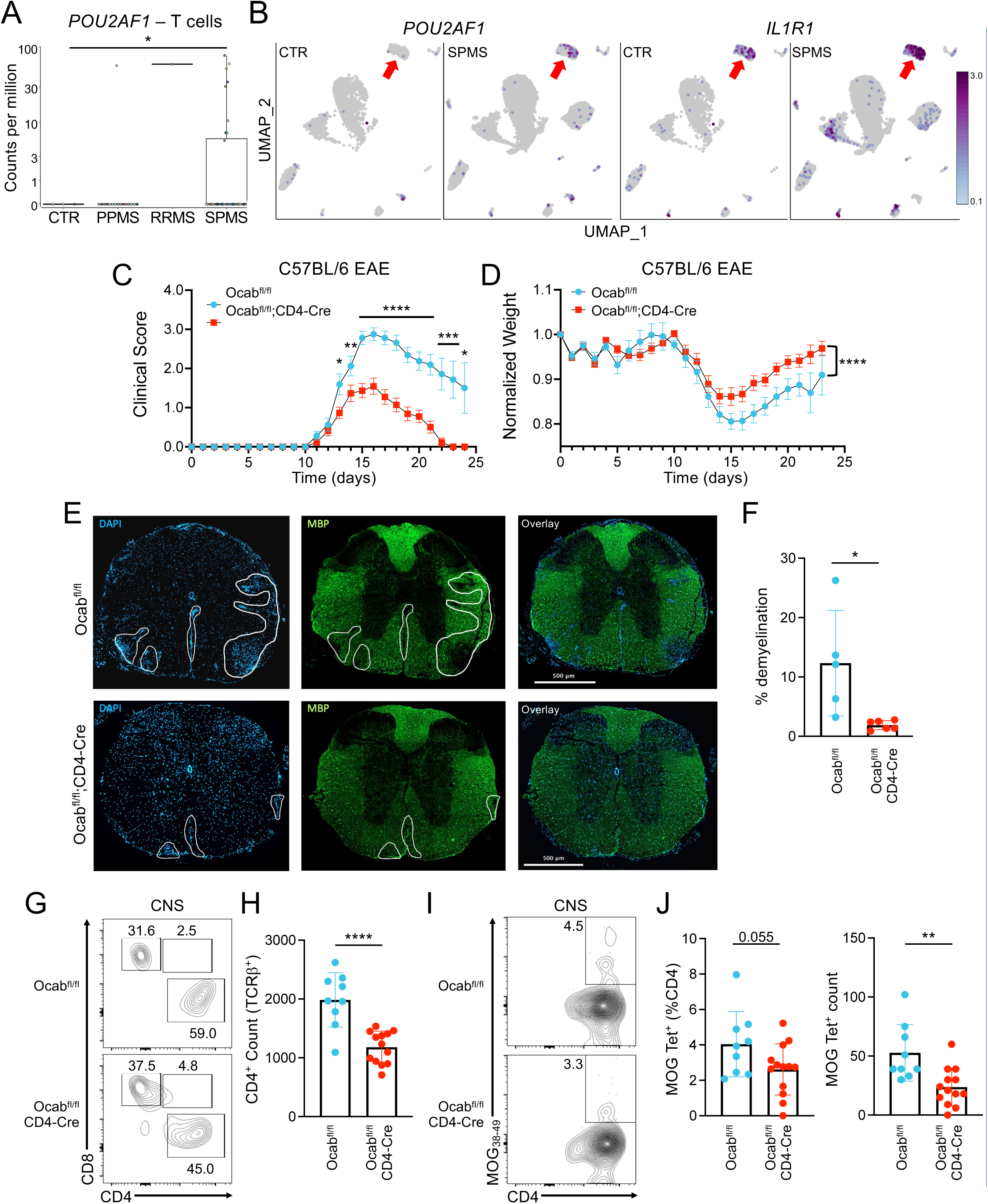
OCA-B loss protects animals from chronic EAE. (**A**) CD4^+^ T cell nuclei from single-nucleus RNAseq data of primary-progressive (PPMS), relapsing-remitting MS (RRMS), and secondary-progressive MS (SPMS) patient lesions and controls (43) were identified and tested in silico for *POU2AF1* expression, which is displayed as counts per million. Significance was ascribed by Welch’s T-test. (**B**) The CD4^+^ T cell nuclei were reclustered and displayed as UMAP feature plots showing *POU2AF1* (left) and *IL1R1* (right) expression in CTR and SPMS brain tissue. Red arrow highlights a likely pathogenic CD4^+^ population expressing high levels of *IL1R1*. (**C**) *Ocab^fl/fl^* (n=16) and *Ocab^fl/fl^*;CD4-Cre (n=22) mice were injected with MOG_35-55_ peptide in CFA and *pertussis* toxin to induce EAE. Clinical scores were evaluated after EAE induction to determine disease progression. (**D**) Animal weights were recorded to evaluate weight loss as a measurement of disease progression. (**E**) Representative DAPI/MBP immunostaining of thoracic spinal cord sections taken from mice 15 days after EAE induction. Areas of demyelination are marked by decreased MBP staining (green) and often coincide with increased cellular infiltration (blue). (**F**) Quantification of demyelination in *Ocab^fl/fl^* (n=5) and *Ocab^fl/fl^*;CD4-Cre mice (n=6). (**G**) Brain and spinal cords were isolated on day 15 of EAE and analyzed by spectral cytometry. Representative flow cytometry plots of CD45^+^TCRβ^+^ CNS infiltrating T cells showing frequency of CD4^+^ and CD8^+^ T cells. (**H**) Quantification of the number of CNS infiltrating CD4^+^ T cells. *Ocab^fl/fl^* (n=9) and *Ocab^fl/fl^*;CD4-Cre (n=13). (**I**) Representative plots showing frequency of MOG_38-49_ tetramer-positive CD4^+^ T cells. (**J**) Quantification of the frequency and count of MOG_38-49_ tetramer-positive CD4^+^ T cells. *Ocab^fl/fl^* (n=9) and *Ocab^fl/fl^*;CD4-Cre (n=13). For clinical scores and normalized weights, values represent mean ±SEM. All other values represent mean ±SD.

### OCA-B promotes chronic EAE in C57BL/6 mice

To understand the impact of OCA-B in T cell mediated autoimmunity, we utilized an *Ocab* conditional allele crossed to the CD4-Cre driver, which efficiently deletes OCA-B in T cells (32). We induced EAE in C57BL/6 female *Ocab^fl/fl^*;CD4-Cre mice and littermate *Ocab^fl/fl^* controls through subcutaneous (s.c.) injection of myelin oligodendrocyte glycoprotein (MOG)_35-55_ peptide in Complete Freund’s Adjuvant (CFA) followed by intraperitoneal (i.p.) injections of *Bordetella pertussis* toxin. *Ocab^fl/fl^*;CD4-Cre mice showed reduced chronic EAE clinical scores and weight loss compared to *Ocab^fl/fl^* (Figure 1C-D). Spinal cords were taken at peak disease (day 15) for immunofluorescence and histological analysis. Myelin basic protein (MBP) immunofluorescence showed a significant decrease in demyelinated white matter in spinal sections from Ocab^fl/fl^;CD4-Cre mice compared to littermate Ocab^fl/fl^ controls while DAPI showed simultaneously decreased immune cell infiltration (Figure 1E-F). Ocab^fl/fl^;CD4-Cre mice also showed a significant decrease in demyelination as measured by luxol fast blue (LFB) in combination with hematoxylin and eosin (H&E) staining (Supplemental Figure 2A-B). Flow cytometric profiling at peak disease showed decreased CD4^+^ T cell CNS infiltration using *Ocab^fl/fl^*;CD4-Cre mice, however no differences in other cell populations, anergic T cells (CD4^+^FR4^+^CD73^+^), or in the relative frequencies of the proinflammatory cytokines IFNy, IL-17, GM-CSF, or CD44 were observed (Figure 1G-H and Supplemental Figure 2C-H). We studied MOG-specific CD4^+^ T cells using MHC class II (I-A^b^) MOG_38-49_ tetramers. At peak disease, *Ocab^fl/fl^*;CD4-Cre mice showed a trending reduction in MOG tetramer-specific cell frequency (*p*=0.055) and a significant reduction in numbers (*p*=0.003) compared to controls (Figure 1I-J). These results indicate that the protection conferred by OCA-B loss in T cells is marked by modest decreases in MOG-specific CD4^+^ T cells within the CNS, resulting in fewer pathogenic T cells driving demyelination.

### OCA-B is essential for EAE mediated by adoptive transfer of Th1- and Th17-polarized T cells

MS is driven by autoreactive CD4^+^ Th1 and Th17 cells within the CNS that recruit CD8^+^ T cells, macrophages, microglia and neutrophils, which together mediate white-matter damage (45). To investigate the impact of OCA-B on autoimmune Th1 and Th17 populations, we used adoptive transfer EAE models with female *Ocab*^fl/fl^;CD4-Cre and *Ocab*^fl/fl^ littermate controls. Mice were primed with MOG_35-55_ in CFA for 14 days. Following MOG priming, CD4^+^ T cells were purified from the lymph nodes of primed experimental or control donors (Supplemental Figure 3A). No differences between genotypes in CD4^+^ T cell viability, count, or effector cytokine (IFNγ and IL-17) expression were observed (Supplemental Figure 3B-E).

To investigate if OCA-B expression impacts Th1 or Th17 mediated adoptive transfer EAE, CD4^+^ T cells purified from donors were polarized in vitro into Th1 and Th17 cells as previously described (46, 47). 3×10^6^ Th1-or Th17-polarized CD4^+^ cells were transferred into naïve age-matched wild-type male C57BL/6 recipients by i.p. injection. EAE onset in mice receiving either Th1-or Th17-polarized control cells was rapid and severe (Figure 2A-B and Supplemental Figure 3F-G). In contrast to control cells, and in contrast to the mild protection observed with OCA-B deficiency in standard C57BL/6 EAE, mice adoptively transferred with T cells lacking OCA-B were almost completely protected from clinical disease regardless of their polarization state (Figure 2A-and Supplemental Figure 3F-G). Interestingly, the ability to initially polarize CD4^+^ T cells into Th1 and Th17 effectors in vitro was only minimally affected by OCA-B loss (Supplemental Figure 3H-J), whereas cells taken from the CNS of engrafted recipient mice at day 15 not only showed decreased CNS infiltration but also reduced IL-17 expression (Figure 2C-F). This effect appeared to be specific to IL-17, as no significant difference in IFNγ was observed (Supplemental Figure 3K). These results indicate a central role for OCA-B in promoting robust neuroinflammatory T cell responses upon antigen reencounter in adoptive transfer EAE models.

**Figure 2.**
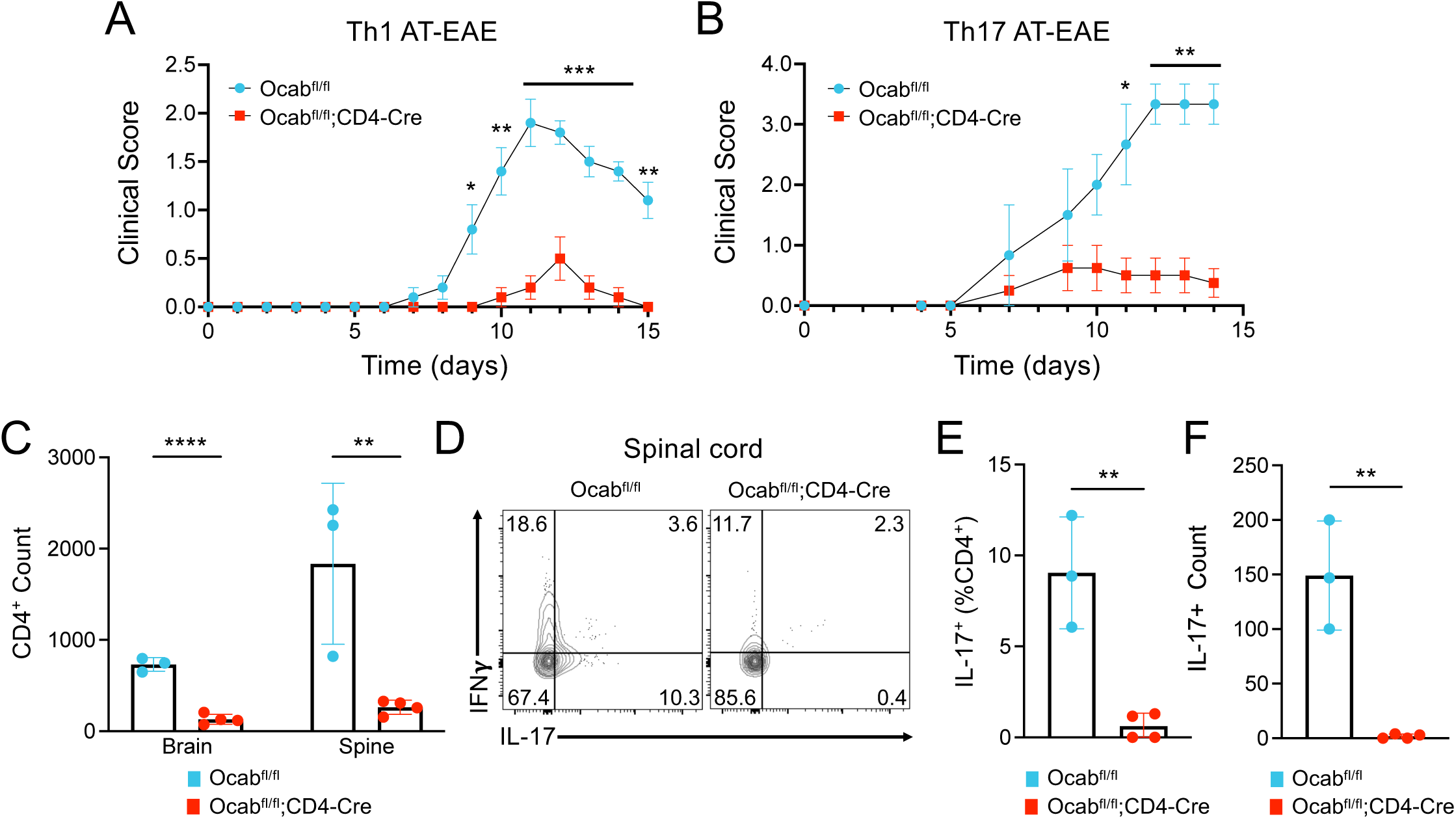
OCA-B promotes Th1 and Th17 adoptive transfer EAE through recall response. (**A,B**) *Ocab^fl/fl^*and *Ocab^fl/fl^*;CD4-Cre mice were primed with MOG_35-55_ peptide in CFA for 10-14 days. Cells from the spleens and lymph nodes of primed mice were isolated and cultured for 4 days in Th1 or Th17 polarizing conditions. 3.0×10^6^ Th1-or Th17-polarized cells were injected i.p. into C57BL/6 wildtype recipient mice (Th1: *Ocab^fl/fl^* n=5, *Ocab^fl/fl^*;CD4-Cre n=5) (Th17: *Ocab^fl/fl^* n=3, *Ocab^fl/fl^*;CD4-Cre n=4). Clinical scores were assessed for 14-15 days. (**C**) At Th17 adoptive transfer EAE endpoint, brains and spinal cords were analyzed by flow cytometry. Quantification of brain- and spinal cord-infiltrating CD4^+^ T cells following Th17 adoptive transfer. (**D**) Representative flow cytometry plots showing the frequency of IFNγ and IL-17 expressing CD4^+^ T cells within the spines of Th17 recipient mice. (**E**) Quantification of IL-17 frequency in spinal cord infiltrating CD4^+^ T cells. (**F**) Quantification of the total count of IL-17^+^ CD4^+^ T cells within the spinal cord of recipient mice. For clinical scores, values represent mean ±SEM. All other values represent mean ±SD.

### OCA-B is dispensable for neurotropic viral clearance

To investigate if T cell OCA-B deletion impacts the ability to mount immune responses and control CNS viral infection, male *Ocab*^fl/fl^;CD4-Cre and *Ocab*^fl/fl^ littermate controls were intracranially inoculated with 1500 plaque-forming units (PFU) of the John H. Muller strain of murine hepatitis virus (JHMV), a β-coronavirus adapted to infect glial cells (astrocytes, microglia and oligodendrocytes) in the mouse brain and spinal cord. Inoculation with JHMV results in acute encephalomyelitis and chronic demyelination similar to autoantigen-driven EAE. Disease severity was determined by evaluating clinical scores over 21 days post-infection (dpi). *Ocab*^fl/fl^;CD4-Cre animals showed negligible differences in clinical scores compared to *Ocab*^fl/fl^ littermates (Figure 3A). Spinal cords were taken on 12 and 21 dpi for immunofluorescence and histological analysis to investigate differences in demyelination. MBP immunofluorescence at 12 dpi showed similar demyelination and immune infiltration between Ocab^fl/fl^ and Ocab^fl/fl^;CD4-Cre spinal sections (Figure 3B-C). Similarly, LFB histology of brain tissue at 21 dpi showed no significant differences in demyelination between OCA-B deficient and control infected animals (Supplemental Figure 4A-B). Flow cytometric analysis of brain tissue showed no significant differences in infiltrating CD4^+^ T cell numbers or IFNγ expression at 7, 12, or 21 dpi (Figure 3D-F, and Supplemental Figure 4C). Similarly, no differences in either total CD8^+^ T cell numbers, or the percentage or numbers of IFNγ-expressing CD8^+^ T cells was observed (Supplemental Figure 4D-F). Brain hemisphere homogenates were collected on 7, 12 and 21 dpi to measure viral titers and clearance. *Ocab*^fl/fl^;CD4-Cre and *Ocab*^fl/fl^ littermate control animals showed similar titers at dpi 7 and 12, and virus was undetectable at day 21 regardless of genotype (Figure 3G). This result suggests that OCA-B is dispensable for baseline immune responses to and clearance of neurotropic viruses.

**Figure 3.**
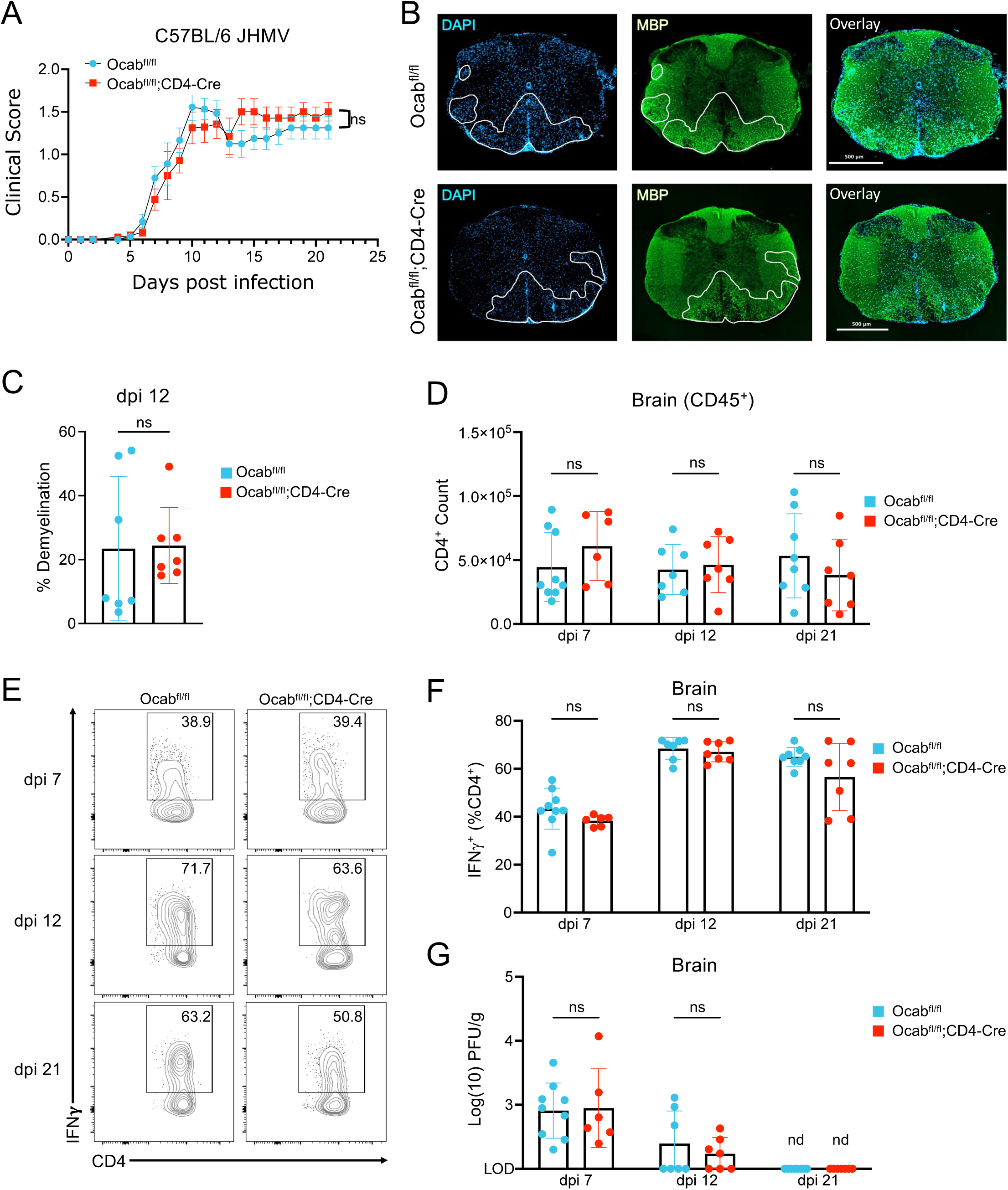
OCA-B is dispensable for T cell response to CNS infection with a neurotropic virus. (**A**) *Ocab^fl/fl^* (n=19) and *Ocab^fl/fl^*;CD4-Cre (n=17) mice were injected intracranially with 1500 PFU JHMV. Clinical scores were recorded for 21 days post infection (dpi). (**B**) Representative DAPI/MBP immunofluorescence staining of spinal cord sections at 12 dpi. Demyelinated areas are marked by decreased MBP staining (green) and often coincide with increased cellular infiltration (blue). (**C**) Quantification of % demyelination in spinal cord sections. (n=7 per group). Half brains were taken at 7, 12, and 21 dpi for flow cytometric and viral titer analysis. (**D**) Quantification of CD4^+^ T cell counts within half brain at 7, 12, 21 dpi. *Ocab^fl/fl^* (n=7-9 per time point) and *Ocab^fl/fl^*;CD4-Cre (n=6-7 per time point). (**E**) Representative plots showing the frequency of IFNγ expressing CD4^+^ T cells 7, 12, and 21 dpi. (**F**) Quantification of frequency of IFNγ expressing CD4^+^ T cells. *Ocab^fl/fl^* (n=7-9 per time point) and *Ocab^fl/fl^*;CD4-Cre (n=6-7 per time point). (**G**) Viral titers of half-brains at dpi 7, 12, and 21 (n.d., not detected). *Ocab^fl/fl^* (n=7-9 per time point) and *Ocab^fl/fl^*;CD4-Cre (n=6-7 per time point). These data represent the combined results of two independent experiments. For clinical scores, values represent mean ±SEM. All other values represent mean ±SD.

### OCA-B expressing CD4^+^ T cells display pathogenic stem-like properties and preferentially transfer demyelinating disease

We used a recently described OCA-B-mCherry reporter mouse (31) to evaluate OCA-B expression within CNS infiltrating T cells. EAE was induced in female C57BL/6 homozygous OCA-B-mCherry reporter mice with MOG_35-55_, and CNS-infiltrating CD4^+^ T cells assessed by flow cytometry at peak clinical disease. mCherry-positive T cells showed increased CD44 and CD62L co-expression, indicative of increased memory-like cells in the OCA-B expressing population (Figure 4A). The association of OCA-B expression with a memory-like profile was observed in the brain, spine and draining cervical lymph nodes (Figure 4B). Interestingly, the frequency of CD44^+^CD62L^-^CD4^+^ T cells was elevated within the OCA-B expressing population specifically within the cervical lymph nodes (Supplemental Figure 5A).

**Figure 4.**
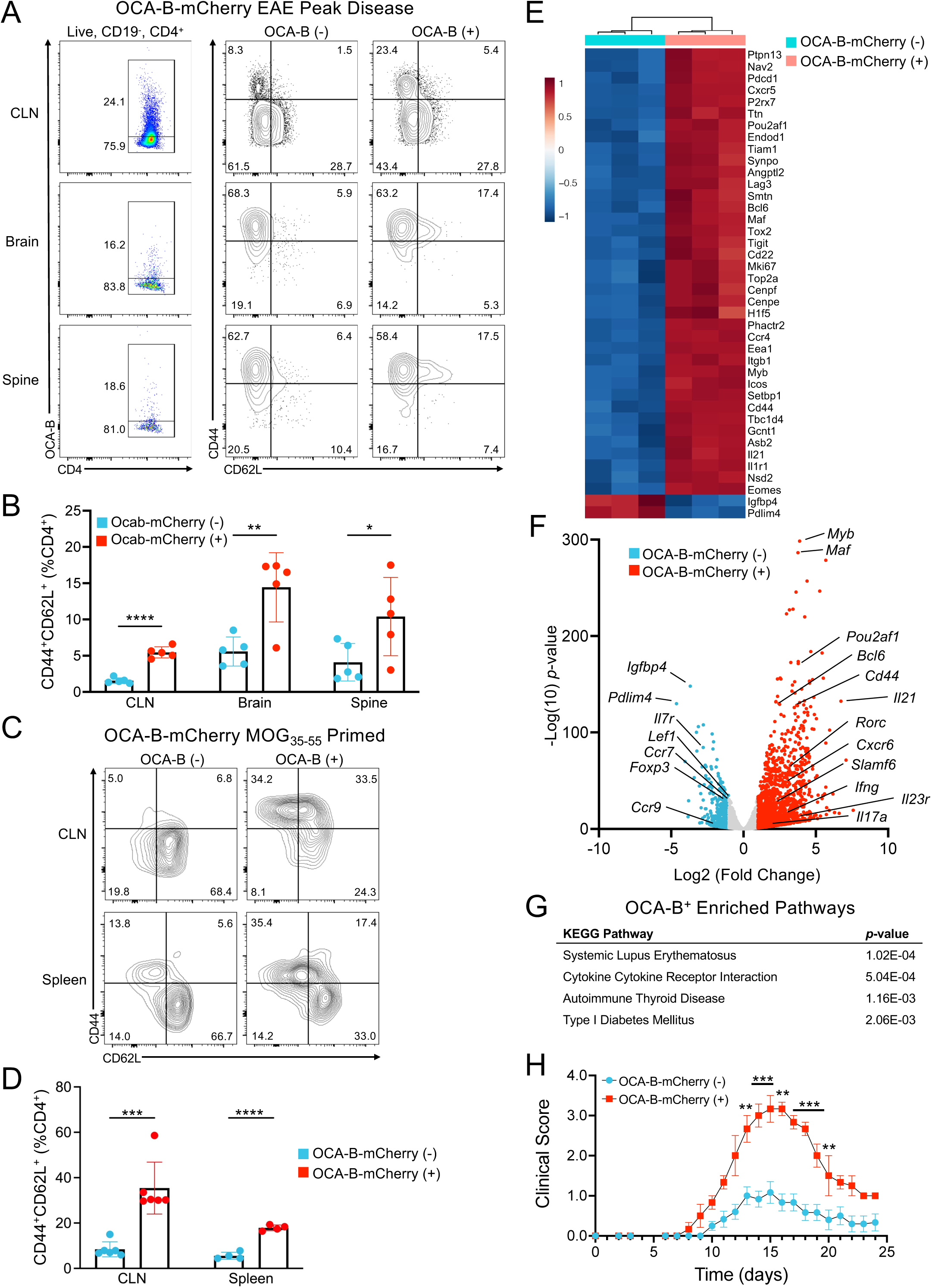
OCA-B expression marks encephalitogenic stem-like CD4^+^ T cells. (**A**) OCA-B-mCherry mice were injected with MOG_35-55_ peptide in CFA and *pertussis* toxin to induce EAE. Cervical lymph nodes (CLNs), brains, and spines were isolated at peak disease (day 15) and analyzed by flow cytometry for mCherry, CD44 and CD62L expression. (**B**) CD44^+^ and CD62L^+^ expressing CD4^+^ T cells were quantified based on mCherry expression. Mean values are shown from n=5 biological replicates. (**C**) OCA-B-mCherry mice were injected with MOG_35-55_ peptide in CFA, and after 14 days CD4^+^ T cells within CLNs and spleens were profiled by flow cytometry for mCherry, CD44, and CD62L expression. (**D**) Quantification of CD44 and CD62L expression of OCA-B negative and positive CD4^+^ T cells from MOG-primed OCA-B-mCherry mice. n=4-6 biological replicates from two combined replicate experiments are shown. (**E**) Heatmap showing relative expression of the top 40 differentially expressed genes from OCA-B positive and negative CD4^+^ T cells from bulk RNA sequencing of OCA-B positive and negative CD4^+^ T cells from MOG_35-55_ primed OCA-B-mCherry reporter mice. (**F**) Volcano plot of differentially expressed genes between OCA-B positive and negative T cells (*p*-value < 0.05 and Log2 fold change > 1 are colored). (**G**) Gene Ontology (GO) terms associated with OCA-B positive and negative groups. (**H**) EAE clinical scores of C57BL/6 wild-type mice following passive transfer of OCA-B-positive or-negative Th1 polarized CD4^+^ T cells. Clinical score values represent mean ±SEM. All other values represent mean ±SD.

Memory-like CD4^+^ T cells have been implicated as major orchestrators of autoimmune demyelination in mice and humans (9–12, 14, 15, 48). To see if this memory-like profile of OCA-B expressing CD4^+^ T cells is specific to CNS infiltrating cells, we performed flow cytometry using draining lymph node and splenic T cells from MOG-primed OCA-B-mCherry reporter mice. Similar to OCA-B^+^ CNS infiltrating cells, OCA-B expressing CD4^+^ T cells taken from the cervical lymph node and spleen showed increased CD44 and CD62L co-expression (Figure 4C-D).

We next evaluated OCA-B-mCherry positive and negative populations by RNA sequencing (RNA-seq) of CD4^+^ T cells isolated from the lymph nodes of 14-day MOG-primed mice. CD4^+^ T cells were sorted based on mCherry expression, as well as lack of CD8, lack of CD19, expression of CD4 and viability (Supplemental Figure 5B). ∼1500 genes were upregulated and ∼350 downregulated in OCA-B expressing compared to OCA-B negative cells (Supplemental Table 1). Upregulated genes in OCA-B expressing cells included *Cxcr6*, *Slamf6*, *Il23r*, *Rorc*, *Ifng*, *Bcl6*, and *Il17a* while downregulated genes included *Foxp3*, *Il7r* and *Ccr9* (Figure 4E-F). Expression of *Tcf7* (which encodes TCF1) was high within both OCA-B positive and negative populations (Supplemental Table 1). Gene ontology (GO) and pathway analysis of upregulated genes show enrichment for terms associated with autoimmune diseases including lupus, autoimmune thyroid disease, and type-1 diabetes (Figure 4G, and Supplemental Table 2). To further assess how OCA-B may transcriptionally regulate these differentially regulated genes we analyzed our previously published Oct1/OCA-B ChIPseq dataset (20). After realignment to the *mm39* reference genome, Oct1 and OCA-B co-localized peaks were observed both near the transcription start site as well as within gene body of *Tcf7*, *Bcl6*, and *Bach2* (Supplemental Figure 5C). These findings indicate that OCA-B likely directly regulates at least a subset of the differentially expressed genes identified in gene knockout experiments.

To determine if OCA-B expression can be used to prospectively identify viable pathogenic CD4^+^ T cell populations, we used OCA-B-mCherry reporter mice as donors in passive transfer EAE. 11-week-old male and female littermates homozygous for the OCA-B-mCherry reporter were primed for 14 days with MOG_35-55_ in CFA as previously described (47). 9×10^5^ mCherry negative and positive cells were cultured in Th1 polarizing conditions and subsequently transferred into age-matched male C57BL/6 recipients as previously described (14). Disease progression was measured by clinical score and normalized weight. Recipients of OCA-B-expressing, mCherry-positive cells showed increased clinical disease severity and weight loss compared to mCherry-negative control recipients (Figure 4H and Supplemental Figure 5D). These findings indicate that OCA-B marks CD4^+^ T cells with pathogenic stem-like gene expression and can be used to prospectively identify peripheral CD4^+^ T cells with enhanced encephalitogenic properties.

### OCA-B promotes neuroinflammation in relapsing-remitting EAE models via control of stem/memory-like T cell populations

In contrast to the chronic form of EAE that develops in C57BL/6 mouse models, EAE on the NOD background exhibits a relapsing-remitting pattern that more closely mimics human MS (42). We induced EAE in 15-18 week old female NOD.*Ocab*^fl/fl^;CD4-Cre and littermate control NOD.*Ocab*^fl/fl^ mice (32) with MOG_35-55_. Clinical scores and weights were recorded for 47 days to observe initial disease onset, relapse and relapse recovery. No significant differences in clinical scores were observed between groups at disease onset (day 10-19), however following the primary remission, NOD.*Ocab*^fl/fl^;CD4-Cre animals were almost completely protected against disease relapse (Figure 5A). Individual NOD mouse clinical scores are shown in Supplemental Figure 6. Normalized weights of animals also reflected the protective effect of OCA-B loss in T cells at relapse (Figure 5B).

**Figure 5.**
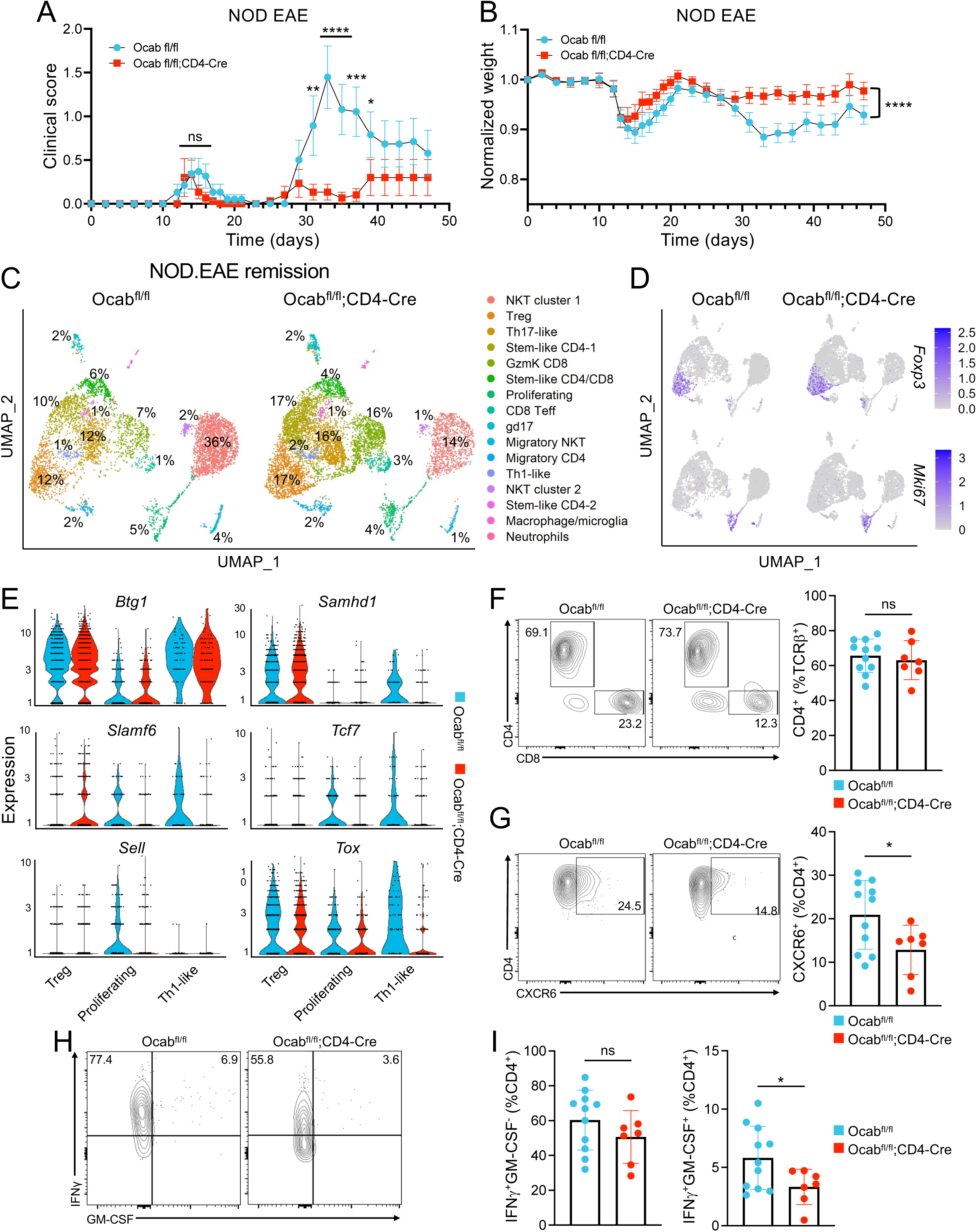
OCA-B promotes relapsing-remitting EAE through stem-like CD4^+^ T cells. (**A**) NOD.*Ocab^fl/fl^* (n=19) and NOD.*Ocab^fl/fl^*;CD4-Cre (n=15) mice were injected with MOG_35-55_ peptide in CFA and *pertussis* toxin to induce EAE. Clinical scores of animals representing relapsing-remitting disease progression are shown as a function of time. (**B**) Animal weights were recorded to determine weight loss throughout initial disease and relapse. (**C**) Singe cell RNA sequencing was performed on pooled CD3͛^+^ cells isolated from the brain and spine of NOD.*Ocab^fl/fl^* (n=3) and NOD.*Ocab^fl/fl^*;CD4-Cre (n=4) 24 days after EAE induction (remission timepoint). Cell populations were plotted in a UMAP using the Seurat R package and percentages are shown for each cluster. Clusters were identified through differential gene expression analysis. (**D**) Feature plots comparing the expression of *Foxp3* and *Mki67* amongst clusters. (**E**) Violin plots showing the expression of *Btg1*, *Samhd1*, *Slamf6*, *Tcf7, Sell* and *Tox* within the Treg, proliferating, and Th1-like clusters. (**F**) CNS cells isolated at remission timepoint (day 24) were analyzed by spectral cytometry. Representative flow cytometry plots and quantification showing the frequency of CD4^+^ T cells between control and experimental groups. (**G**) Representative flow cytometry plots and quantification showing the frequency of CXCR6 expressing CD4^+^ T cells. (**H**) Representative flow cytometry plots showing the frequency of CD4^+^ T cells expressing IFNγ and GM-CSF. (**I**) Quantification of CD4^+^ T cells expressing IFNγ and GM-CSF. For clinical scores, data represent mean ±SEM. All other data represents mean ±SD.

To understand differences in CNS-infiltrating T cells underlying relapse protection at a population and transcriptional level, we performed single-cell RNA-seq (scRNA-seq) of CD3ε^+^ T cell isolates from CNS (brain + spinal cord) at remission (day 24) and peak relapse (day 33). Cells from 3-6 NOD.*Ocab*^fl/fl^;CD4-Cre and NOD.*Ocab*^fl/fl^ littermate controls were collected and combined for analysis. 7799 (remission) and 9224 (relapse) *Ocab* knockout, and 6009 (remission) and 7525 (relapse) control cells passed filtering and were used for analysis, with an average per cell read depth ranging from 90,000 to 190,000. For remission, UMAP clustering using both genotypes revealed 16 clusters corresponding to a broad T cell repertoire consisting of natural-killer T (NKT) cells (the largest cluster), γδ T cells with a Th17-like gene signature (gd17), and different CD4^+^ and CD8^+^ subsets (Figure 5C and Supplemental Figure 7A). Noteworthy clusters based on mRNA expression included proliferating (*Ki67*, *Pcna*, *Mcm2*), Th1-like (*Tbx21*, *Ifng*, *Tnf*, *Il12rb2*) and Treg (*Foxp3*, *Ctla4*, Supplemental Figure 7A-B and Supplemental Table 3). Feature plots for *Foxp3* and *Mki67* are shown in Figure 5D. Additional feature plots for *Cxcr6*, *Il23r* and *Bach2* are shown in Supplemental Figure 7C. *Bach2* was notably poorly expressed in all clusters, in contrast to relapse timepoints (below). Notably, there was a large decrease in NKT cell numbers in knockout mice (36% control vs. 14% experimental). Within this cluster, changes in gene expression between control and conditional knockout groups were minimal. Most NKT cells from both groups showed a limited inflammatory cytokine response, consistent with the lack of clinical disease at remission. The decrease in the large NKT population in the experimental group proportionally increases the percentage contribution of the other clusters, and therefore similar or decreased percentages of these cells identifies reduced populations which correlate with protection from relapse, e.g. proliferating T cells. Other clusters such as Tregs increased disproportionately. Differential gene expression analysis of key clusters (Treg, Proliferating, and Th1-like) revealed increased expression of *Slamf6* in Tregs lacking OCA-B, while OCA-B deficient proliferating and Th1-like cells showed decreased expression of genes associated with quiescence (*Btg1* and *Samhd1*), memory/stemness (*Slamf6*, *Tcf7*, and *Sell*) and pathogenicity (*Tcf7* and *Tox* co-expression, Figure 5E). Spectral cytometry showed no significant difference in the frequency of CNS infiltrating CD4^+^ T cells between groups at remission (Figure 5F). However, OCA-B deficient CD4^+^ T cells did show a significant reduction in CXCR6 and IFNγ/GM-CSF co-expression, indicative of reduced Th17 pathogenicity (Figure 5G-I). These results indicate that loss of OCA-B in T cells selectively reduces proliferation and differentiation of pathogenic stem/memory-like CD4^+^ T cells during disease remission.

At relapse, multiple CNS-infiltrating stem-like and effector CD4^+^ and CD8^+^ control T cell populations were present (Figure 6A and Supplemental Figure 8A-B, Supplemental Table 4). TCR sequencing revealed an expansion of a subset of TCR clones in the OCA-B deficient CD8 clusters while most other clusters were highly polyclonal (Figure 6B). Comparing the 10 most expanded TCR clonotypes between knockouts and controls, there were several clonotypes that were shared but differentially expanded (Supplemental Figure 8C and Supplemental Table 5). A “Ccr9 stem-like” cluster expressing the gut homing chemokine receptor *Ccr9*, as well as *Ccr7*, *Bach2*, and *Tcf7* notably appeared at relapse (Figure 6C and Supplemental Table 4). TCR clonotype analysis indicated that this population was highly polyclonal (Figure 6B). Additionally, a “Stem-like CD4-2” cluster expressing *Tcf7*, Ccr7, *Bach2, Cd44* and low-level *Cxcr6*, and a prominent “Th17-like” cluster expressing *Tcf7, Cd44*, *Il1r1, Lgals3*, and higher levels of *Cxcr6* was identified (Figure 6C). This latter cluster likely contains pathogenic effector cells.

**Figure 6.**
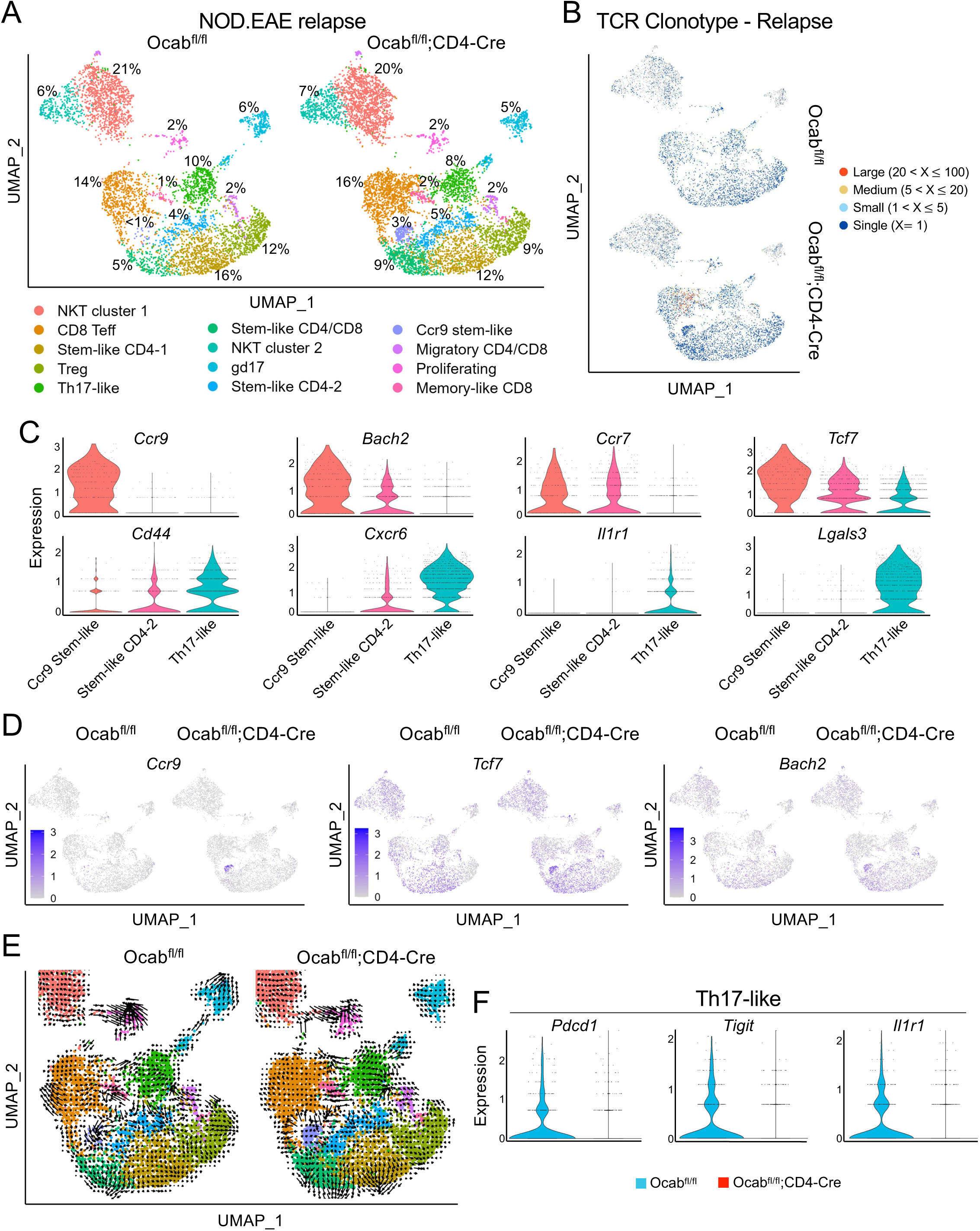
OCA-B promotes disease relapse through control of pathogenic stem-like Th17 differentiation. (**A**) Singe cell RNA and TCR sequencing were performed on pooled CD3͛^+^ cells isolated from the brain and spine of NOD.*Ocab^fl/fl^* (n=5) and NOD.*Ocab^fl/fl^*;CD4-Cre (n=6) 33 days after EAE induction (relapse timepoint). Cell populations were plotted in a UMAP using the Seurat R package and percentages are shown for each cluster. Clusters were identified through differential gene expression analysis. (**B**) UMAP TCR clonotype expansion between *Ocab^fl/fl^* and *Ocab^fl/fl^*;CD4-Cre amongst clusters. (**C**) Violin plots showing expression of *Ccr9*, *Bach2*, *Ccr7*, *Tcf7*, *Cd44*, *Cxcr6*, *Il1r1* and *Lgals3* in the Ccr9 Stem-like CD4, Stem-like CD4-2, and Th17-like clusters. (**D**) Feature plots comparing the expression of *Ccr9*, *Tcf7*, and *Bach2* amongst clusters. (E) UMAPs showing RNA velocity of spliced and unspliced transcripts by experimental condition. (F) Violin plots showing expression of *Pdcd1*, *Tigit*, and *Il1r1* in the Th17-like cluster between control and experimental groups.

In the absence of OCA-B, NKT cell populations were normalized compared to the remission timepoint, and several clusters showed major changes in the percentage of cells including most prominently within the Ccr9-stem-like cluster which was much more prevalent (Figure 6D). Feature plots comparing OCA-B T cell deficient mice to littermate controls for *Ccr9* and the transcription factors *Tcf7* and *Bach2* are shown in Figure 6D. The cells were also marked by the expression of *Cd4*, *Tox, Ccr7, Ccr4* and limited expression of *Sell* (Supplemental Figure 9A).

RNA velocity analysis of spliced and unspliced transcripts (49) identified clusters of T cells, most bearing static gene expression profiles consistent with terminal differentiation, but some with differentiating or transitional gene expression (Figure 6E). Notably, OCA-B knockout cells showed increased transcriptional stasis within the Ccr9-stem-like and proliferating clusters, and a decreased transcriptional directionality of Th17 cells towards gd17 cells, the latter of which express high levels of pathogenic markers including *Il1r1*, *Cxcr6*, and *Lgals3* (Supplemental Figure 6A). Pseudotime analysis identified a transcriptional capacity in the control *Ccr9* expressing population to differentiate into pathogenic Th17-like cells, whereas cells lacking OCA-B were limited in this capacity, possibly resulting in the accumulation of these cells (Supplemental Figure 9B). These changes culminated in the Th17-like effector knockout cells expressing reduced pathogenicity-associated genes such as *Pdcd1*, *Tigit* and *Il1r1* (Figure 6F). Together these results suggest OCA-B mediates the transition of non-pathogenic stem-like cells to pathogenic Th17 cells which ultimately drive disease relapse.

## Discussion

The development of therapies that inhibit autoimmunity while minimizing impacts on normal immune function remains a critical unmet goal in the field. In this study, we show that expression of mRNA encoding the transcriptional co-regulator OCA-B is elevated in CD4^+^ T cells from SPMS brain tissue and from effector memory-like peripheral blood CD4^+^ T cells from RRMS patients. We show that OCA-B promotes the pathogenic maturation of T cells and establishment of CNS autoimmunity, in particular episodes of relapse. Loss of OCA-B attenuates these populations in chronic EAE, in adoptive transfer EAE, and in NOD.EAE during remission. During relapse, OCA-B T cell knockout mice are almost completely protected, and accumulate a population of CD4^+^ T cells expressing *Ccr9*, *Bach2*, and *Tcf7*. OCA-B expression also marks viable pathogenic stem/memory-like CD4^+^ T cells that preferentially transfer EAE. Furthermore, the effects of modulating OCA-B are selective, as responses to infection with a neurotropic coronavirus that generates similar clinical symptoms but lacks an autoantigen were largely unaffected.

OCA-B is a member of a small family of transcription co-regulators that also includes OCA-T1, OCA-T2 and IκBε (50, 51). It becomes induced upon CD4^+^ T cell activation docks with Oct1 to directly regulate immunomodulatory genes such as *Il2*, *Ifng*, *Il17a*, Tbx21 (T-bet) and *Csf2* (*Gmcsf*). Rather than functioning as a primary activator of these genes, it is required for their robust expression upon antigen reencounter (20, 32). Notably, unlike transcription factors such as NF-AT, AP-1 and NF-κB, Oct1 and OCA-B are dispensable for normal primary T cell responses (20). These findings predict important roles for OCA-B in CD4^+^ T cells specifically in situations of antigen reencounter, which is a necessary feature of autoimmunity. Conditional T cell knockout of Oct1 reduces EAE severity, with reduced proinflammatory cytokine expression and increased anergy (21). Deletion of Oct1 in T cells was dispensable for neurotropic viral clearance (21), however the ubiquitous expression of Oct1 in adult tissue limits its translational potential. In contrast, OCA-B expression is largely confined to B and T cells, and whole-body knockout mice are viable and fertile (52). OCA-B is highly expressed in B cells and required for germinal center formation (52, 53). It is expressed in CD4^+^ T cells at levels 50-100-fold less than in B cells (32, 54, 55). This suggests that with proper therapeutic dosing, OCA-B could be inhibited selectively in T cells with minimal impact to the B cell compartment. Short-term administration of an OCA-B peptide mimic that inhibits OCA-B’s downstream effector functions reverses spontaneous type-1 diabetes in NOD mice (32). However, toxicity would limit the peptide mimic’s efficacy, including in EAE models. Further research is required for the development of safe and effective OCA-B inhibitors.

OCA-B whole body knockout animals are partially protected from EAE (41), however, the known activity of OCA-B in B cells made it unclear if the protection is T cell-derived. We show that OCA-B promotes EAE pathogenesis through T cells. Opportunistic infections and reactivation of latent CNS-tropic viruses are significant barriers for MS therapeutics that focus on global immune suppression as a therapeutic mechanism. When OCA-B T cell conditional knockout mice are challenged with the neurotropic coronavirus JHMV, there is little difference in their ability to mount immune responses and control virus replication. These data are consistent with previous findings that OCA-B loss in T cells preserves lymphocytic choriomeningitis virus (LCMV) acute viral infection response, instead selectively impacting memory recall responses (20, 31). This finding provides a potential therapeutic window in which targeting the low levels of OCA-B in T cells could be used to blunt autoimmunity while sparing baseline immune responses.

Unlike the significant but modest protection observed in chronic C57BL/6 EAE, OCA-B loss in adoptive transfer EAE almost completely protects recipient animals from disease. This may indicate that adoptive transfer EAE models involve more autoantigen re-exposures, as transferred T cells must re-encounter antigen in a naïve host before promoting disease. Protection was observed regardless of whether donor cells were Th1-or Th17-polarized, indicating that OCA-B functions through a mechanism that parallels rather than enforces the Th1/Th17 paradigm. We therefore sought other mechanisms that might explain why OCA-B T cell knockouts are so strongly protected.

Autoimmune pathogenesis and maintenance has been increasingly associated with stem-or progenitor-like T cell subpopulations (56, 57). Recent findings indicate an emerging role for stem/memory-like T cells in promoting autoimmunity (48, 58–60). For example, myelin-specific CD4^+^ T cells in MS patients show signs of a previously activated, memory-like phenotype compared to healthy controls (9, 10, 12). Furthermore, memory-like Th1 and Th17 cells are increased in the peripheral blood of MS patients and display increased pathogenicity compared to conventional Th17 Cells (61). Similarly, in EAE models, peripheral central memory-like CD4^+^CD44^hi^CD62L^hi^ T cells confer increased disease severity upon transfer (14). Both Oct1 and OCA-B promote the functionality of long-lived CD4^+^ memory T cells following antigenic rechallenge (31, 62). An OCA-B-mCherry reporter mouse allele, which effectively labels memory progenitor CD4^+^ T cells during acute infection (31), also labels CD4^+^ T cells that preferentially confer EAE. OCA-B^hi^ cells taken from the CNS at peak EAE selectively express stem/memory markers. Further profiling OCA-B^hi^ CD4^+^ T cells suggests that OCA-B promotes the transition of non-pathogenic stem-like cells to a pathogenic state associated with markers such as *Cxcr6*, *Slamf6*, *Ifng*, and *Il17a*. Elevated *Pou2af1* (*Ocab*) transcripts are also found in pathogenic Th17 cell populations (63). The connection between OCA-B expression and pathogenic memory/stem-like T cells likely has not been appreciated before because OCA-B is expressed at levels too low to be reproducibly captured with many current single-cell RNA-seq technologies.

Relapsing-remitting EAE models have been developed using the Swiss Jim Lambert (SJL) and autoimmune prone non-obese diabetic (NOD) backgrounds (42, 64). In NOD mice, MOG_35-55_ EAE has been described as relapsing-remitting with minimal initial disease that increases in severity upon relapse (65, 66). Using this model with the conditional *Pou2af1* allele backcrossed to the NOD background (32), we found that OCA-B T cell loss minimally affects initial clinical presentation, but instead protects animals specifically at relapse. The finding that OCA-B loss in conventional C57BL/6 EAE results in partial protection intermediate between the initial and relapse phases of NOD.EAE suggests that chronic C57BL/6 EAE may not model either initial presentation or relapse accurately, but instead represents a superposition of the two. Nevertheless there are caveats associated with the NOD.EAE model, including poor characterization relative to the C57BL/6 and SJL models, relatively weak clinical scores, lack of temporal synchronic between replicate animals after the first relapse, and lack of underlying progression (42, 67). Although we find that *POU2AF1* transcripts are elevated in T cells from lesions of patients with secondary-progressive MS, more work is necessary for confirm selectivity for OCA-B in promoting patient relapse and underlying progression.

Single-cell RNA expression profiling at remission identifies a significant loss of *Slamf6*, *Tcf7* (encoding TCF1) and *Sell* from proliferative stem-like CD4^+^ T cells in the OCA-B deficient condition. While the majority of research on TCF1-expressing stem-like T cells focuses on CD8^+^ populations with the capacity for self-renewal and differentiation into effector cells, analogous descriptions have been made for CD4^+^ populations (48, 68, 69). Indeed, autoimmune CD4^+^ T cells expressing these memory/stem-like markers have been reported as an encephalitogenic reservoir that play a key role in EAE pathogenicity (48). In the same study, bulk RNA sequencing associated OCA-B expression within this stem-like reservoir (48).

Single-cell RNA-seq at relapse identified multiple emergent populations of effector T cells compared to remission. Notably, there was also an accumulation of a unique population of CD4^+^ cells in OCA-B T cell deficient mice. These cells are conspicuously marked by expression not only of *Ccr4*, but also of *Ccr9*, which encodes a gut-homing chemokine receptor. CCR9^+^ memory-like CD4^+^ T cells have been linked to MS and EAE protection (70). The cells also express *Bach2*, a transcription regulator which was recently associated with non-pathogenic immunomodulatory Th17-polarized cells in the context of C57BL/6 EAE (71). *Bach2* expression was largely concentrated in the “CCR9 stem-like” cluster, which is more numerous in the knockout condition. Consistently, in bulk RNA-seq *Bach2* mRNA was upregulated in OCA-B^-^ cells (*p*<10^-13^), although with a small (+1.74) fold change (Supplemental Table 1). Similarly, recent work associates *Pou2af1* (*Ocab*) with pathogenic Th17/Th1 cells compared to the non-pathogenic *Bach2* expressing cells (71). In B cells, OCA-B is known to regulate *Bach2* (72), however the interplay between OCA-B and Bach2 and how this influences the pathogenic maturation of Th17 cells merits additional research. Velocity and pseudotime analysis indicates that these cells are normally transcriptionally inclined to progress towards an highly pathogenic Th17 population, but that progression stalls in mice whose T cells lack OCA-B. Consistently, the Th17 effector population in OCA-B T cell knockout mice lacks expression of key genes such as *Il1r1* and *Pdcd1*, both of which are associated with pathogenicity in CNS-infiltrating Th17 cells^48,64^. The cumulative alterations in these populations in OCA-B knockouts likely explains why these mice are strongly protected against disease relapse. In conclusion, this study indicates a prominent role for OCA-B in driving stem-like CD4^+^ T cell populations towards a pathogenic state while playing a minimal role in acute antiviral responses to a CNS infection.

## Methods

### Sex as a biological variable

The objective of this study was to determine the role of the transcriptional regulator OCA-B in demyelinating disease. OCA-B T cell conditional knockout mice and OCA-B reporter mice were utilized in multiple mouse demyelinating disease models to evaluate the impact of T cell OCA-B expression on clinical disease manifestation and the phenotypes of neuroinflammatory cells, in particular T cells. Both male and female animals were used for experiments except for active EAE experiments where only female animals were used as these are the better model spinal cord demyelination and inflammation.

### Analysis of publicly available human control and MS single nuclei RNAseq dataset

Initial evaluation of *POU2AF1* expression in T cells was conducted in the interactive web browser created by the authors of the dataset available at https://malhotralab.shinyapps.io/MS_broad/ (43). Briefly, *POU2AF1* differential gene expression was filtered to show T cell gene expression by diagnosis with a minimal threshold cutoff of 10 cells per sample. Cleaned aligned matrix files were then downloaded from https://zenodo.org/records/8338963 and subsequently processed by using the 10X Genomics CellRanger pipeline and further analyzed using the 10X Genomics Loupe Browser 7 software. 19,906 CD4^+^ cells were then subclustered and evaluated for *POU2AF1* and *IL1R1* gene expression by diagnosis.

### Chronic C57BL/6 EAE

11-12 week-old female C57BL/6 littermate animals were used for all experiments. For adoptive transfer experiments using the OCA-B-mCherry reporter allele (31), 11 week-old females were used for chronic EAE and 9 week-old males were used as MOG_35-55_ primed adoptive transfer donors. EAE was induced on day 0 by two 100 μL s.c. injections into both hind flanks of 0.5 μmol/mL MOG_35-55_ peptide (University of Utah Peptide Core) emulsified in complete Freud’s adjuvant (CFA), comprised of incomplete Freund’s adjuvant (Cat# 77145;ThermoFisher) together with 4 mg/mL heat-killed *M. tuberculosis* (Cat#DF3114338, Fisher Scientific). Animals were given two 400 ng i.p. injections of *B. pertussis* toxin (Cat#181, List Biological Laboratories) on days 0 and 2.

### Clinical Scoring

EAE clinical scores were determined using a 0-5 scale with 0.5 scoring intervals: 0.5, tip of tail is limp or tail muscle strain observed; 1, complete limp tail; 1.5, complete limp tail and at least one hind limb consistently falls through wire rack when animal is dropped; 2, complete limp tail and one hind leg dragging without movement beyond hip; 2.5, complete limp tail and dragging of both hind limbs without movement beyond hip; 3, complete hind limb paralysis; 3.5, complete hind limb paralysis with flat laying hind quarters and hump in upper torso; 4, complete hind limb and partial forelimb paralysis, minimal movement around cage; 4.5, complete hind limb and partial forelimb paralysis, no movement around cage; 5, seizure or death. Animals scoring a ≥4 for more than 48 hr were euthanized and scored 5 for the remainder of the experiment. JHMV clinical disease was evaluated using previously described 0-4 point scale (74).

### Immunofluorescence and Histology

Spinal columns were isolated from C57BL/6 Ocab^fl/fl^ and Ocab^fl/fl^;CD4-Cre animals after EAE or JHMV induced demyelination and fixed overnight in in 4% paraformaldehyde at 4C. Spinal cords were subsequently isolated and cryoprotected in 30% sucrose for 3-5 days. Thoracic and lumbar sections were embedded in optimum cutting temperature (OCT) compound (ThermoFisher Scientific) and frozen and stored at −80C. 8 μm spinal sections were cut by cryostat and stained with Luxol fast blue (Cat#212170250; ThermoFisher) along with hematoxylin (Cat#8495; ENG Scientific) and eosin (Cat#E511-25; Fisher Chemical). Demyelination was assessed by dividing the area of demyelinated white matter by total white matter as previously described (75, 76). For immunofluorescence, spinal cord sections were blocked in 5% normal goat serum (NGS; Cat#005-000-121, Jackson ImmunoReseach) and 0.1% Tween-20 for 1 hr at room temperature. Sections were stained overnight at 4°C with primary antibody rabbit anti-MBP (1:1000; Cat#PA1050, Boster), followed by secondary antibody goat anti-rabbit conjugated with Alexa Fluor 488 (1:750; Cat#A11008, Invitrogen) for 2 hr at room temperature. Primary and secondary antibodies were diluted in 1% bovine serum albumin (BSA) and 0.1% Tween-20. Slides were cover-slipped using Fluoromount-G with DAPI (Cat#00495952, Invitrogen). Sections were imaged with a Zeiss Axioscan 7 Microscope Slide Scanner using a 20× objective.

### Flow cytometry and spectral cytometry

Spinal cords and brains were harvested at peak disease or specified timepoint and mechanically dissociated by passing through a 100 μm cell strainer. A 30/70% Percoll gradient (Cat#17544501, Cytiva) was used to enrich mononuclear cells. Cells from collected spleens and lymph nodes (cervical, brachial, axillary and inguinal) were passed through a 70 μm strainer and erythrocytes lysed by ammonium-chloride-potassium lysis buffer (150 mM NH_4_Cl, 10 mM KHCO_3_, 0.1 mM Na_2_EDTA). For intracellular staining, cells were cultured in complete media (RPMI 1640, 10% FBS, 1% Pen/Strep, Glutamax) supplemented with 1 μL/mL brefeldin A (Cat#555029, Golgiplug, BD), 50 ng/mL phorbol myristate acetate (Cat# P1585; Sigma-Aldrich) and 1 μg/mL ionomycin (Cat#I0634, Sigma-Aldrich) for 4 hr. Mouse I-A(b) MOG_38-49_ tetramers (GWYRSPFSRVVH) (77, 78) and control I-A(b) Human CLIP tetramers (PVSKMRMATPLLMQA) conjugated to APC or PE were synthesized by the National Institutes of Health tetramer core facility. Cells were stained with control CLIP or MOG_38-49_ tetramers for 1 hr at 37C prior to live/dead, surface, and intracellular staining. Cultured cells were fixed using cytofix/cytoperm (Cat# 554714; BD) according to manufacturer’s protocol and stained for intracellular cytokines in perm/wash buffer. Antibodies used for flow cytometry include: CD45-Percp(30F11, Cat#103129, Biolegend), CD11b-APC/Cy7(M1/70, Cat#101225, Biolegend), CD19-BUV661(ID3, Cat#612971, BD), CD19-FITC(103/CD19, Cat#152404, Biolegend), NK1.1-PE-Cy5(PK136, Cat#108715, Biolegend), CD3e-BV605(17A2, Cat#100237, Biolegend), CD3e-FITC(145-2C11, Cat#100305, Biolegend), TCRb-BV570(H57-597, Cat#109231, Biolegend), CD4-BV711(RM4-5, Cat#100557, Biolegend), CD4-BUV395(GK1.5, Cat#565974, BD), CD8a-APC(53-6.7, Cat#100712, Biolegend), CD8a-BUV737(53-6.7, Cat#612759, BD), I-A/I-E-BUV805(M5/144.15.2, Cat#748844, BD), CD62L-PE-Cy7(MEL-14, Cat#25-0621-82, eBioscience), CD44-Percp-Cy5.5(IM7, Cat#45-0441-80, eBioscience), Ly-108-BUV563(13G3, Cat#7741436, BD), FR4-BUV496(12A5, Cat#750415, BD), CD73-V450(Ty/23, Cat#561544, BD), IL-23R-BV421(12B2B64, Cat#150907, Biolegend), CXCR6-BV711(SA051D1, Cat#151111, Biolegend), PD-1-APC/Fire810(29F.1A12, Cat#135251, Biolegend), IFNγ-PE-Cy7(XMG1.2, Cat#25-7311-82, eBioscience), IFNγ-APC(XMG1.2, Cat#17-7311-81, eBioscience), IL-17a-BV605(TC11-18H10.1, Cat#506927, Biolegend), IL-17a-BV650(TC11-18H10.1, Cat#506929, Biolegend), TNFa-FITC(MP6-XT22, Cat#506304, Biolegend), and GMCSF-PE/Dazzle594(MP1-22E9, Cat#505421, BioLegend). Flow cytometry samples were profiled using a BD Fortessa (BD Biosciences, Figure 2, 3, 4) or an Aurora spectral flow cytometer (Cytek, Fig 1 and 5). FlowJo software (BD Biosciences) was used for data analysis.

### Pre-polarization profiling and mixed adoptive transfer

10 week-old C57BL/6 *Ocab^fl/fl^* and *Ocab^fl/fl^;CD4-Cre* littermates were immunized with MOG_35-55_ in CFA as described above. After 14 days, spleen and lymph node (cervical, axillary, brachial and inguinal) cells were isolated and cultured in complete media (RPMI 1640, 10% FBS, 1% Pen/Strep, Glutamax) supplemented with 1 μL/mL brefeldin A (Cat#555029; Golgiplug, BD), 50 ng/mL phorbol myristate acetate (Cat#P1585; Sigma-Aldrich) and 1 μg/mL ionomycin (Cat#I0634, Sigma-Aldrich) for 4 hr. Cells were stained for surface markers, fixed using the Foxp3/Transcription Factor Staining Buffer Kit (Cat#00-5523-00, ThermoFisher), and stained for intracellular markers (IFNγ and IL-17).

### Th1 and Th17 polarization and adoptive transfer C57BL/6 EAE

10-12 week-old female C57BL/6 littermate animals were used for all experiments. Donor animals were immunized with MOG_35-55_ in CFA as above. 10-14 days post-immunization, spleen and lymph node (cervical, axillary, brachial and inguinal) cells were collected and placed into mixed culture with MOG peptide and cytokines at 5.0×10^6^ cells/mL. For Th1 conditions, complete media was supplemented with 50 μg/mL MOG_35-55_, 6 ng/mL rmIL-6 (Cat#216-16, Peprotech) and 2 ng/mL rmIFNγ (Cat#315-05, Peprotech). For Th17 conditions, complete media was supplemented with 50 μg/mL MOG_35-55_, 8 ng/mL rmIL-23 (Cat#1887-ML-010, R&D Systems), 10 ng/mL rmIL-1α (Cat#211-11A, Peprotech), and 10 μg/mL anti-IFNγ (Cat#BE0055, BioXcell XMG1.2). After 4 days of respective culture, CD4^+^ T cells were purified using negative magnetic selection and resuspended in PBS at 1.5×10^6^ cells/mL. 100 μL cell suspension was injected i.p. into age-matched male C57BL/6 recipients (∼13 week-old at time of disease induction). Clinical disease was scored for 15 days.

### JHMV

*Ocab^fl/fl^* and *Ocab^fl/fl^*;CD4-Cre C57BL/6 mice were anesthetized using isoflurane. Mice were intracranially (i.c) injected with 1500 plaque-forming units (PFU) of JHMV (strain V34) suspended in 30 μL Hank’s balanced salt solution (ThermoFisher). Clinical disease was assessed for 21 days using a previously described scale (74). Mice were sacrificed at days 7, 12, and 21 post-infection to assess viral titers within brain homogenates using a plaque assay previously described (79). Spinal cords were taken on day 12 and 21 post-infection to assess demyelination by DAPI/MBP and luxol fast blue and hematoxylin/eosin histology respectively.

### OCA-B-mCherry adoptive transfer EAE

OCA-B-mCherry reporter allele animals (31) were primed for 14 days with MOG_35-55_ in CFA as described above. After priming, spleen and lymph nodes were collected and CD4^+^ T cells were purified by magnetic negative selection (Miltenyi Bio kit and LS columns). Non-CD4^+^ T cells were removed from the LS column and placed on ice for subsequent in vitro culture. Purified CD4^+^ T cells were separated by OCA-B-mCherry expression (PE-Texas Red) using flow automated cell sorting (FACS) on a BD Aria cell sorter. OCA-B-mCherry positive and negative CD4^+^ T cells were placed into mixed culture in complete medium with the previously purified non-CD4^+^ T cells at a 1:10 ratio (CD4^+^ to non-CD4^+^) at 1.0×10^6^ cells/mL. The culture was supplemented with 20 μg/mL MOG_35-55_ and 0.5 ng/mL IL-12p70 (Cat#210-12, Peprotech). After two days of culture, CD4^+^ T cells were again purified through negative magnetic selection. Purified CD4^+^ T cells from OCA-B-mCherry positive and negative cultures were resuspended in PBS at 8.8×10^6^ cells/mL. 100 μL cell suspension was injected i.p. into age-matched male C57BL/6 recipients. Clinical disease was assessed for 24 days.

### Bulk RNA-seq

OCA-B-mCherry reporter mice (n=9) were primed for 14 days with MOG_35-55_ in CFA as described above. After priming, lymph nodes (cervical, axillary, brachial and inguinal) were isolated and mCherry positive and negative CD4^+^ T cells were collected by FACS. Total RNA, three groups (pooled 3 mice each), was isolated using the Quick RNA micro kit (Cat# R1050; Zymo Research) with DNase treatment (Cat#79254, Qiagen). RNA concentration was measured with a Qubit RNA HS Assay Kit (Cat#Q32855, Fisher Scientific). RNA quality was evaluated with an RNA ScreenTape Assay (Cat#5067-5579, 5067-5580, Agilent Technologies). Total RNA samples (5-500 ng) were hybridized with NEBNext rRNA Depletion Kit v2 (Cat#E7400, New England Biolabs) to diminish rRNA from the samples. Stranded RNA sequencing libraries were prepared as described using the NEBNext Ultra II Directional RNA Library Prep Kit for Illumina (Cat#E7760L; New England Biolabs). Purified libraries were qualified on an 4150 TapeStation (Agilent Technologies) using a D1000 ScreenTape assay (Cat#5567-5582, 5067-5583, Agilent Technologies). The molarity of adapter-modified molecules was defined by quantitative PCR using the Kapa Biosystems Kapa Library Quant Kit (Cat#KK4824, Roche). Individual libraries were normalized to 5 nM. NovaSeq 150×150 bp sequencing libraries were chemically denatured and applied to an Illumina NovaSeq flow cell using the XP workflow (20043131). Following transfer to an Illumina NovaSeq 6000 instrument, 150×150 cycle paired-end sequencing was performed using a S4 reagent Kit v1.5 (Cat#20028312, Illumina). 8-11 million reads were generated per sample and were aligned to the *GRCm38* mouse reference genome. Differential gene expression and pathway analysis were conducted using the DESeq2 and Enrichr R packages.

### NOD-EAE

15-18 week-old female littermate *Ocab^fl/fl^* and *Ocab^fl/fl^*;CD4-Cre animals backcrossed to a NOD strain background (32) were used for all experiments. Diabetic mice (blood glucose >150 mg/dL) were excluded from the analysis. EAE was induced by two 100 μL s.c. injections into both hind flanks of 0.5 μmol/mL MOG_35-55_ peptide emulsified in incomplete Freuds adjuvant (Cat#77145, ThermoFisher) supplemented with 4 mg/mL heat-killed *M. tuberculosis* (Cat#DF3114338, Fisher Scientific) to form CFA. Animals were given two 400 ng i.p. injections of *Bordetella pertussis* toxin (Cat#181, List Biological Laboratories) on days 0 and 2. Clinical disease was assessed over 47 days using the same 5-point scale described above. Remission timepoints were determined to be between days 22-26 while first relapse was determined to be between days 26-39.

### Single-cell RNA-seq

NOD.*Ocab^fl/fl^* and NOD.*Ocab^fl/fl^*;CD4-Cre animals were induced with EAE and monitored for disease progression. Brains and spinal cords were isolated on day 24 (remission) and day 33 (relapse), and processed for FACS sorting. Viable cells were sorted by CD3͛ using a FACSAria (BD Bioscience). Cells from each condition were isolated from 3-4 mice and combined. Cells were processed using the 5’ 10X Genomics Chromium platform according to the manufacturer’s instructions. Paired-end high-throughput sequencing (125 cycles) was performed using a NovaSeq instrument (Illumina). Sequencing reads were processed by using the 10X Genomics CellRanger pipeline and further analyzed using the Seurat R package. Analysis of cells used a standard filtering protocol removing cells with unique feature counts of >5,000 or <100, as well as cells with >10% mitochondrial counts (indicative of dead cells). No more than 10% of total cells were removed by this process. Cells were subjected to unsupervised hierarchical clustering followed by uniform manifold approximation and projection (UMAP) to visualize clusters with similar gene expression and their relative positions. RNA velocity analysis was performed using velocyto.R version 0.6 to model cell state transitions using the guidelines provided in Manno et al. (49) and the associated repository at https://github.com/velocyto-team/velocyto.R. High dimensional velocity vectors were graphed on the predefined Seurat UMAP projection using localized average grid projections. Pseudotime analysis was conducted using Seurat and Monocle3 packages. Integrated data was divided into subset based on genotype and Monocle3 was used to calculate trajectory, order cells and plot pseudotime for each condition. *Statistics.* Multiple two-tailed student’s t-tests were used to determine statistical differences in EAE clinical scores and all error bars denote ±SEM unless otherwise noted. Two-way ANOVA was used to determine differences in weight loss throughout disease progression. GraphPad Prism software was used for all statistics and graphing. Two-tailed student’s t-tests were used to determine statistical differences in cell populations observed by flow cytometry and all error bars represent ±SD unless otherwise noted. For all figures, ns = *p*>0.05, * = *p*≤0.05, ** = *p*≤0.01, *** = *p*≤0.001, **** = *p*≤0.0001.

### Study approval

All animal experiments were performed in strict accordance with the NIH guide for the Care and Use of Laboratory animals and institutional guidelines for animal care at the University of Utah under approved protocol #00001553.

## Data availability

All RNA sequencing, single-cell sequencing, and single-cell T cell V(D)J sequencing data associated with the findings of this study have been deposited to NCBI’s GEO database under accession code GSE243727. Data values for all graphs and values behind any reported means are located in the Supporting Data Values file.

## Author contributions

E.P.H. and D.T. conceived the study. E.P.H. and A.S. performed experiments. E.P.H, A.S., E.M.M. and B.T.S. analyzed data. T.E.L., B.T.S. and D.T. provided critical resources and supervision for the study. E.P.H. and D.T. wrote, reviewed and revised the manuscript.

## Supporting information

Supplemental Figures 1-9 with legends

## Acknowledgements

We thank R.M. O’Connell and M. Bettini for critical reading of the manuscript. We thank J. Marvin and University of Utah Health Sciences Center Flow Cytometry core, and B. Dalley, O. Allen and the High-Throughput Genomics core. Peptides were synthesized by the DNA/Peptide facility. We thank T. Lane for assistance with JHMV constructs, protocols and reagents. This work was supported by grants from the National Institutes of Health/ National Institute of Allergy and Infectious Diseases (R01AI162929 and R01AI100873) and the Praespero Foundation (Canada) to DT, and training grants from the National Institute of Neurological Disorders and Stroke (T32NS115664) to EPH and National Institute of Allergy and Infectious Diseases (T32AI055434) to ARS.

## Supplemental figure legends

**Supplemental Figure 1. Reclustered CD4^+^ nuclei gene expression and analysis of peripheral blood CD4^+^ T effector cell bulk RNAseq.** (**A**) Additional feature plots of reclustered CD4^+^ T cell nuclei from single-nucleus RNAseq data of secondary-progressive MS (SPMS) patient lesions and controls (43) showing cluster gene expression of *CD4*, *CD44*, *LEF1*, *ETS1*, *BCL6*, and *MAF*. (**B**) Quantification of *POU2AF1* counts from bulk RNAseq data of CD4^+^ T effector cells isolated from the peripheral blood of control (HC Teff) and MS (MS Teff) patients (44). All data represent mean ±SD and Welch’s T-test was used to determine statistical significance for data with unequal variance.

**Supplemental Figure 2. Minimal differences in cytokine expressing CD4^+^ T cell, CD8^+^ T cell, B cell, macrophage or microglial cell counts at peak chronic EAE in OCA-B knockouts.** (**A**) Representative spinal cord LFB/H&E staining of thoracic sections taken from mice 15 days after EAE induction. Areas of demyelination are outlined in black and marked by increased loss of LFB (blue) staining and visibility of H&E (pink) within the white matter. (**B**) Quantification of the % demyelination from LFB/H&E histology. (**C**) Quantification of the frequency of CNS infiltrating CD4^+^ T cells between Ocab^fl/fl^ and Ocab^fl/fl^;CD4-Cre mice. (**D**) Quantification of frequency and count of CD8α positive T cells between Ocab^fl/fl^ and Ocab^fl/fl^;CD4-Cre mice. (**E**) Quantification of the number of B cells, macrophages and microglia. (**F**) Representative flow cytometry plots comparing the expression of FR4 and CD73 within CNS infiltrating CD4^+^ T cells from Ocab^fl/fl^ and Ocab^fl/fl^;CD4-Cre mice. (**G**) Quantification of the frequency of CNS infiltrating FR4^+^ CD73^-^ and FR4^+^ CD73^+^ CD4^+^ T cells within Ocab^fl/fl^ and Ocab^fl/fl^;CD4-Cre mice. (**H**) Quantification of frequency and count of IFNγ, IL-17, GM-CSF, and CD44 expressing CD4^+^ T cells from Ocab^fl/fl^ and Ocab^fl/fl^;CD4-Cre mice. All data represent mean ±SD.

**Supplemental Figure 3. Loss of OCA-B does not strongly alter initial Th1 or Th17 polarization in vitro.** (**A**) Experimental schematic for assessing CD4^+^ T cell differences between Ocab^fl/fl^ and Ocab^fl/fl^CD4-Cre prior to *in vitro* Th1 or Th17 polarization. Lymph nodes (cervical, brachial, axillary, and inguinal) were harvested from Ocab^fl/fl^ (n=4) and Ocab^fl/fl^;CD4-Cre (n=4) mice following 14 days of MOG_35-55_/CFA priming and analyzed by flow cytometry. (**B**) Frequency of viable cells isolated from Ocab^fl/fl^ and Ocab^fl/fl^;CD4-Cre mice. (**C**) Total CD4^+^ T cell numbers isolated from Ocab^fl/fl^ and Ocab^fl/fl^;CD4-Cre mice. (**D**) Quantification of the frequency of IFNγ expressing CD4^+^ T cells isolated from Ocab^fl/fl^ and Ocab^fl/fl^;CD4-Cre mice. (**E**) Quantification of the frequency of IL-17 expressing CD4^+^ T cells derived from Ocab^fl/fl^ and Ocab^fl/fl^;CD4-Cre mice. (**F**) Normalized weights of recipient mice following adoptive transfer of Ocab^fl/fl^ or Ocab^fl/fl^;CD4-Cre Th1-polarized MOG-reactive CD4^+^ T cells. (**G**) Normalized weights of recipient mice following adoptive transfer of Ocab^fl/fl^ or Ocab^fl/fl^;CD4-Cre Th17 cells. (**H**) Representative flow cytometry plots showing IFNγ and IL-17 expressing CD4^+^ T cells from Ocab^fl/fl^ (n=5) and Ocab^fl/fl^;CD4-Cre (n=5) mice after Th1 and Th17 *in vitro* polarization. (**I**) Quantification of IFNγ expressing CD4^+^ T cells after Th1 polarization. (**J**) Quantification of IL-17 expressing CD4^+^ T cells after Th17 polarization. (**K**) Quantification of the frequency of IFNγ expressing CD4^+^ T cells isolated from the spinal cords of Th17 adoptive transfer EAE recipient mice. Normalized weight variance is displayed as mean ±SEM. All other data represent mean ±SD.

**Supplemental Figure 4. Minimal changes in immune responses to intracranial JHMV infection in the absence of OCA-B.** (**A**) Representative LFB/H&E histology of spinal cord sections at dpi 21. (**B**) Demyelination quantification of LFB/H&E histology at dpi 21. (**C**) Quantification of the number of IFNγ expressing CD4^+^ T cells at dpi 7, 12 and 21. (**D**) Quantification of the number of CD8^+^ T cells at dpi 7, 12 and 21. (**E**) Quantification of the frequency of IFNγ expressing CD8^+^ T cells at dpi 7, 12 and 21. (**F**) Quantification of the number of IFNγ expressing CD8^+^ T cells at dpi 7, 12 and 21. All data represent mean ±SD.

**Supplemental Figure 5. Sorting OCA-B-mCherry reporter T cells and additional analysis of the effects of transferring OCA-B^hi^ versus OCA-B^lo^ CD4^+^ cells.** (**A**) Quantification of the frequency of CD44^+^ CD62^-^ CD4^+^ T cells within the cervical lymph nodes of OCA-B-mCherry reporter mice at peak EAE by OCA-B-mCherry expression. (**B**) Gating strategy for sorting of OCA-B^+^ and OCA-B^-^ CD4^+^ T cells. (**C**) *mm39* ChIP-seq tracks showing peaks for Oct1 (blue), OCA-B (red), along with vertebrate conservation (black) at *Tcf7*, *Bcl6*, and *Bach2*. (**D**) Normalized weights of recipient mice following adoptive transfer of Ocab mCherry positive or negative Th1 cells. Normalized weight is represented by mean ±SEM. All other data represent mean ±SD.

**Supplemental Figure 6. Individual NOD mouse EAE clinical scores.** (**A**) Clinical scores of individual NOD.Ocab^fl/fl^ mice following EAE induction. (**B**) Clinical scores of individual NOD.Ocab^fl/fl^;CD4-Cre mice following EAE induction.

**Supplemental Figure 7. Additional remission scRNA-seq cluster annotation, heat map of gene enrichments by cluster, and additional feature plots.** (**A**) Violin plots showing cluster gene expression of *CD3e*, *Cd4*, *Cd8a*, *Ncr1*(NKp46), *Cxcr6*, *Tcf7*, and *Tcrg-V6* at EAE remission (**B**) Heatmap showing the top 10 genes expressed by cluster at remission EAE timepoint. (**C**) UMAP feature plots showing expression of *Cxcr6*, *Il23r*, and *Bach2* amongst clusters at remission.

**Supplemental Figure 8. scRNA-seq cluster annotation genes, heat map of NOD.EAE gene enrichments by cluster, and TCR clonotype overlap between conditions at the relapse timepoint.** (**A**) Violin plots showing cluster gene expression of *CD3e*, *Cd4*, *Cd8a*, *Ncr1*(NKp46), *Cxcr6*, *Tcf7*, and *Tcrg-V6* at EAE relapse. (**B**) Heatmap showing the top 10 genes expressed by cluster at relapse EAE timepoint. (**C**) Alluvial plot showing top 10 TCR clonotype gene overlap between Ocab^fl/fl^ and Ocab^fl/fl^;CD4-Cre groups. (**C**) UMAP feature showing the expression of *CD4*, *CD8a, Tox*, *Sell*, *Ccr7*, and *Ccr4* amongst clusters at relapse timepoint.

**Supplemental Figure 9. Example relapse feature plots and pseudotime analysis of relapse single cell RNA seq.** (**A**) UMAP feature showing the expression of *CD4*, *CD8a, Tox*, *Sell*, *Ccr7*, and *Ccr4* amongst clusters at relapse timepoint. (**B**) Pseudotime analysis of relapse single cell from root starting node at or near the *Ccr9*-expressing stem-like cluster.

