## Supplemental Figures 1-9 with legends for "OCA-B promotes pathogenic maturation of stem-like CD4^+^ T cells and autoimmune demyelination"

Supplementary figures

### Supplemental Figure 1

A

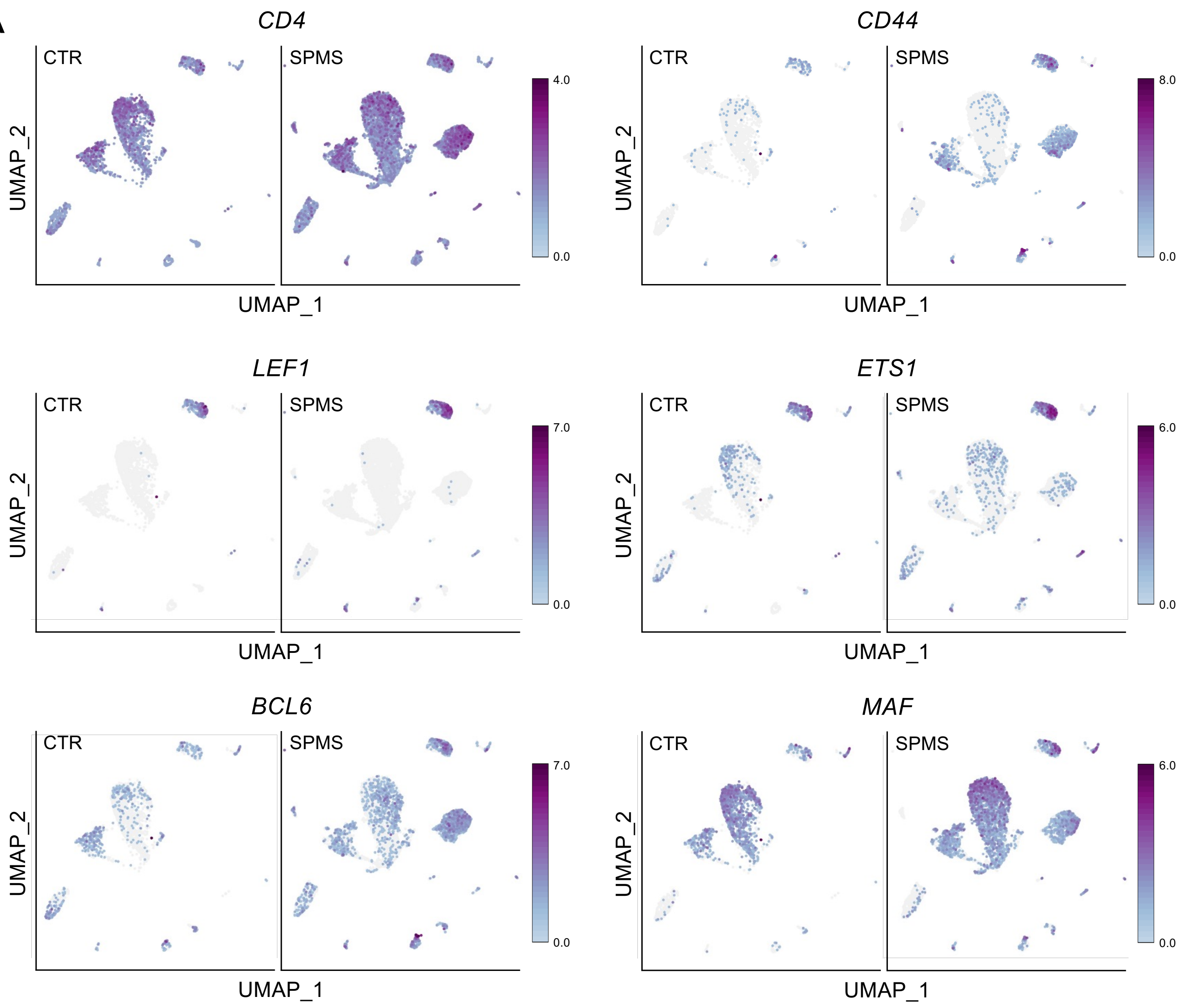


B

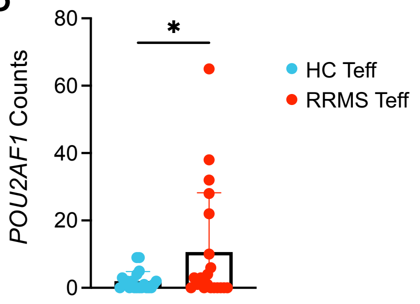

**Supplemental Figure 1. Reclustered CD4$^+$ nuclei gene expression and analysis of peripheral blood CD4$^+$ T effector cell bulk RNAseq .** (**A**) Additional feature plots of reclustered CD4$^+$ T cell nuclei from single-nucleus RNAseq data of secondary-progressive MS (SPMS) patient lesions and controls (McNair et al. 2025 Neuron 113:396) showing cluster gene expression of *CD4*, *CD44*, *LEF1*, *ETS1*, *BCL6*, and *MAF*. (**B**) Quantification of *POU2AF1* counts from bulk RNAseq data of CD4$^+$ T effector cells isolated from the peripheral blood of control (HC Teff) and MS (MS Teff) patients (Sumida et al. 2024 Sci. Trans. Med. 16:eapd1720). All data represent mean ±SD and Welch's T-test was used to determine statistical significance for data with unequal variance.

### Supplemental Figure 2

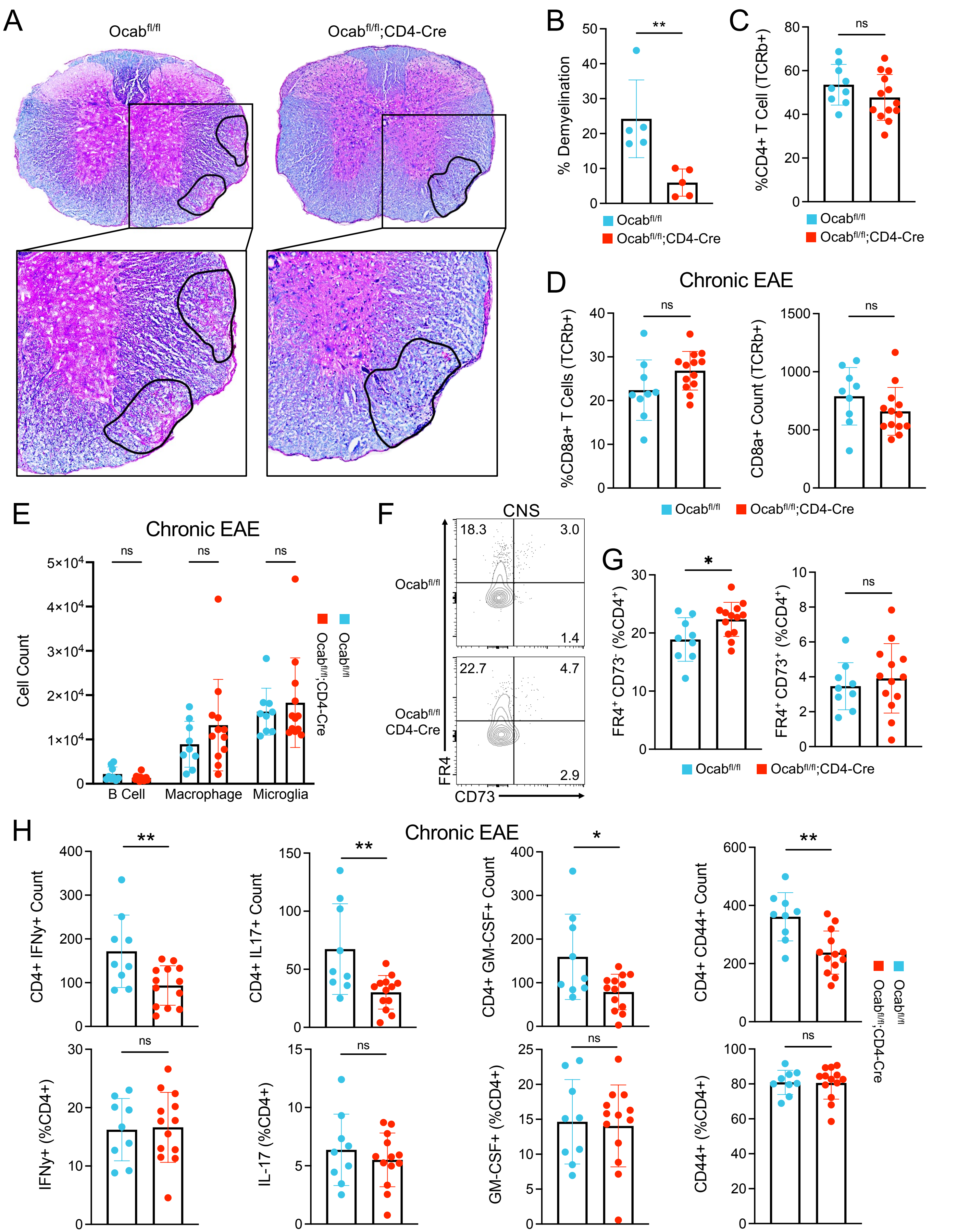

**Supplemental Figure 2. Minimal differences in cytokine expressing $CD4^+$ T cell, $CD8^+$ T cell, B cell, macrophage or microglial cell counts at peak chronic EAE in OCA-B knockouts.** (**A**) Representative spinal cord LFB/H&E staining of thoracic sections taken from mice 15 days after EAE induction. Areas of demyelination are outlined in black and marked by increased loss of LFB (blue) staining and visibility of H&E (pink) within the white matter. (**B**) Quantification of the % demyelination from LFB/H&E histology. (**C**) Quantification of the frequency of CNS infiltrating $CD4^+$ T cells between $Ocab^{fl/fl}$ and $Ocab^{fl/fl}$;CD4-Cre mice. (**D**) Quantification of frequency and count of CD8$\alpha$ positive T cells between $Ocab^{fl/fl}$ and $Ocab^{fl/fl}$;CD4-Cre mice. (**E**) Quantification of the number of B cells, macrophages and microglia. (**F**) Representative flow cytometry plots comparing the expression of FR4 and CD73 within CNS infiltrating $CD4^+$ T cells from $Ocab^{fl/fl}$ and $Ocab^{fl/fl}$;CD4-Cre mice. (**G**) Quantification of the frequency of CNS infiltrating $FR4^+$ $CD73^-$ and $FR4^+$ $CD73^+$ $CD4^+$ T cells within $Ocab^{fl/fl}$ and $Ocab^{fl/fl}$;CD4-Cre mice. (**H**) Quantification of frequency and count of IFN$\gamma$, IL-17, GM-CSF, and CD44 expressing $CD4^+$ T cells from $Ocab^{fl/fl}$ and $Ocab^{fl/fl}$;CD4-Cre mice. All data represent mean $\pm$SD.

### Supplemental Figure 3

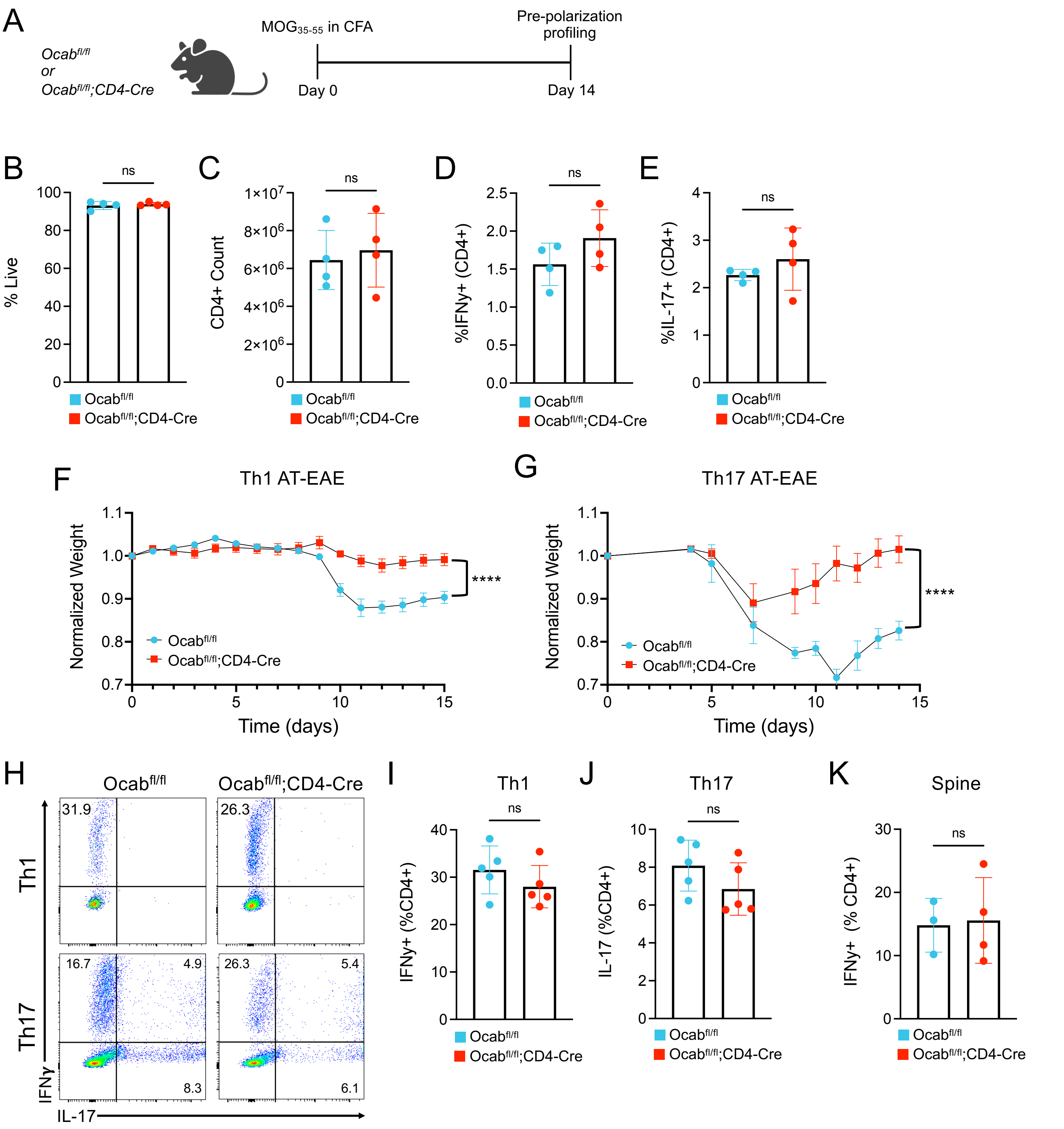

**Supplemental Figure 3. Loss of OCA-B does not strongly alter initial Th1 or Th17 polarization in vitro.** (**A**) Experimental schematic for assessing CD4$^+$ T cell differences between Ocab$^{fl/fl}$ and Ocab$^{fl/fl}$CD4-Cre prior to *in vitro* Th1 or Th17 polarization. Lymph nodes (cervical, brachial, axillary, and inguinal) were harvested from Ocab$^{fl/fl}$ (n=4) and Ocab$^{fl/fl}$;CD4-Cre (n=4) mice following 14 days of MOG$_{35-55}$/CFA priming and analyzed by flow cytometry. (**B**) Frequency of viable cells isolated from Ocab$^{fl/fl}$ and Ocab$^{fl/fl}$;CD4-Cre mice. (**C**) Total CD4$^+$ T cell numbers isolated from Ocab$^{fl/fl}$ and Ocab$^{fl/fl}$;CD4-Cre mice. (**D**) Quantification of the frequency of IFN$\gamma$ expressing CD4$^+$ T cells isolated from Ocab$^{fl/fl}$ and Ocab$^{fl/fl}$;CD4-Cre mice. (**E**) Quantification of the frequency of IL-17 expressing CD4$^+$ T cells derived from Ocab$^{fl/fl}$ and Ocab$^{fl/fl}$;CD4-Cre mice. (**F**) Normalized weights of recipient mice following adoptive transfer of Ocab$^{fl/fl}$ or Ocab$^{fl/fl}$;CD4-Cre Th1-polarized MOG-reactive CD4$^+$ T cells. (**G**) Normalized weights of recipient mice following adoptive transfer of Ocab$^{fl/fl}$ or Ocab$^{fl/fl}$;CD4-Cre Th17 cells. (**H**) Representative flow cytometry plots showing IFN$\gamma$ and IL-17 expressing CD4$^+$ T cells from Ocab$^{fl/fl}$ (n=5) and Ocab$^{fl/fl}$;CD4-Cre (n=5) mice after Th1 and Th17 *in vitro* polarization. (**I**) Quantification of IFN$\gamma$ expressing CD4$^+$ T cells after Th1 polarization. (**J**) Quantification of IL-17 expressing CD4$^+$ T cells after Th17 polarization. (**K**) Quantification of the frequency of IFN$\gamma$ expressing CD4$^+$ T cells isolated from the spinal cords of Th17 adoptive transfer EAE recipient mice. Normalized weight variance is displayed as mean ±SEM. All other data represent mean ±SD.

### Supplemental Figure 4

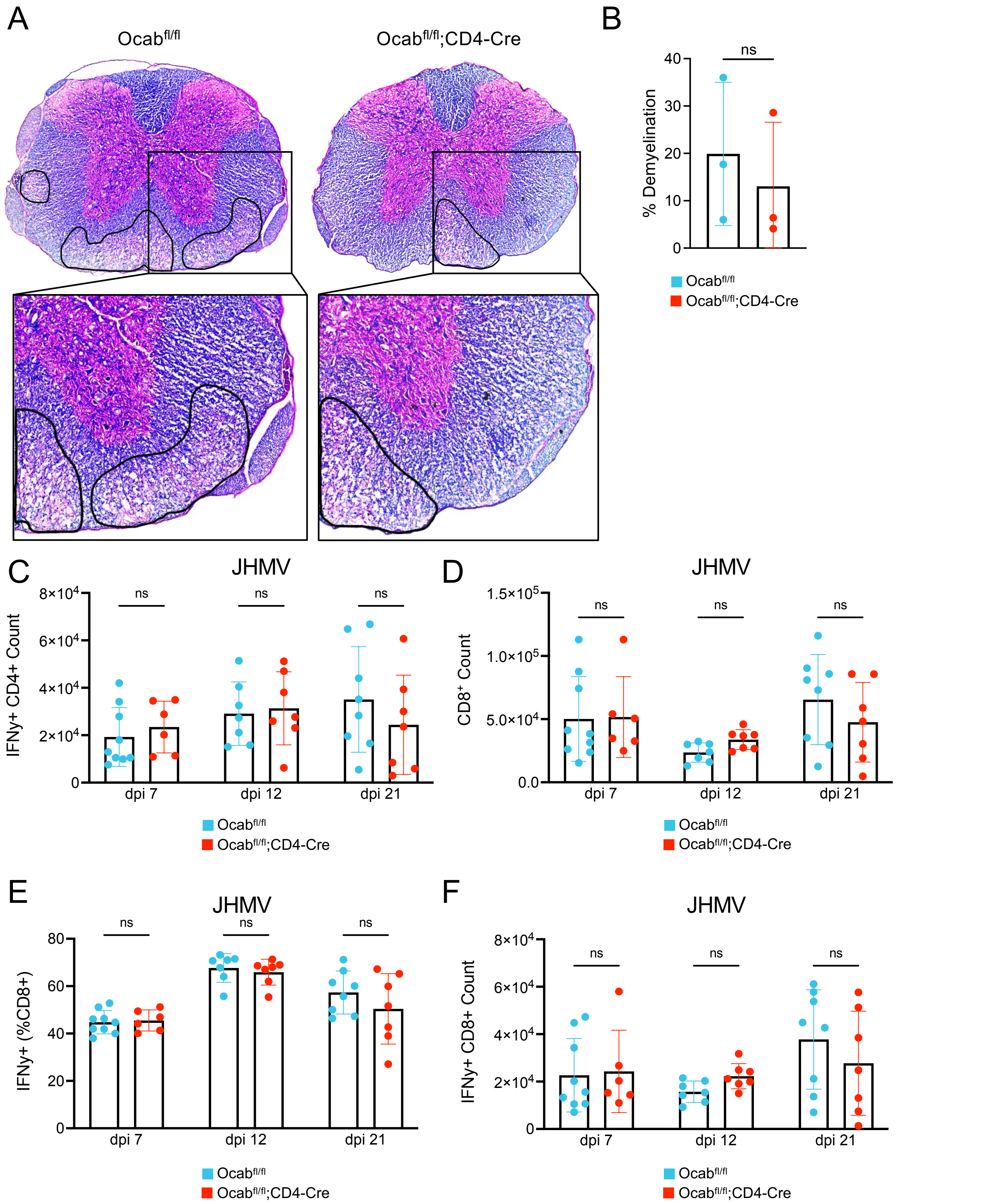

**Supplemental Figure 4. Minimal changes in immune responses to intracranial JHMV infection in the absence of OCA-B.** (**A**) Representative LFB/H&E histology of spinal cord sections at dpi 21. (**B**) Demyelination quantification of LFB/H&E histology at dpi 21. (**C**) Quantification of the number of IFNγ expressing $CD4^+$ T cells at dpi 7, 12 and 21. (**D**) Quantification of the number of $CD8^+$ T cells at dpi 7, 12 and 21. (**E**) Quantification of the frequency of IFNγ expressing $CD8^+$ T cells at dpi 7, 12 and 21. (**F**) Quantification of the number of IFNγ expressing $CD8^+$ T cells at dpi 7, 12 and 21. All data represent mean ±SD.

### Supplemental Figure 5

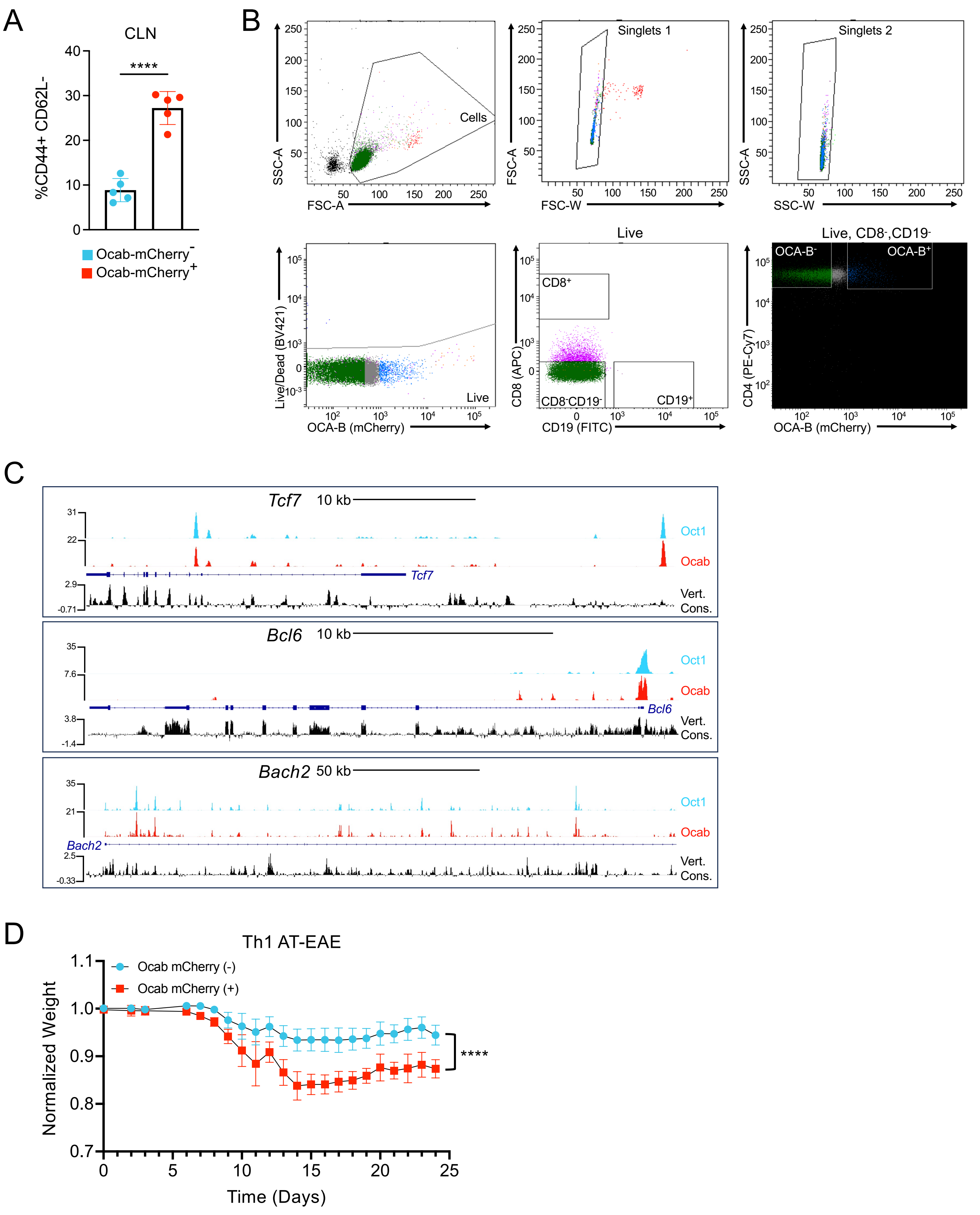

**Supplemental Figure 5. Sorting OCA-B-mCherry reporter T cells and additional analysis of the effects of transferring OCA-B$^{hi}$ versus OCA-B$^{lo}$ CD4$^{+}$ cells.** (**A**) Quantification of the frequency of CD44$^{+}$ CD62$^{-}$ CD4$^{+}$ T cells within the cervical lymph nodes of OCA-B-mCherry reporter mice at peak EAE by OCA-B-mCherry expression. (**B**) Gating strategy for sorting of OCA-B$^{+}$ and OCA-B$^{-}$ CD4$^{+}$ T cells. (**C**) *mm39* ChIP-seq tracks showing peaks for Oct1 (blue), OCA-B (red), along with vertebrate conservation (black) at *Tcf7*, *Bcl6*, and *Bach2*. (**D**) Normalized weights of recipient mice following adoptive transfer of Ocab mCherry positive or negative Th1 cells. Normalized weight is represented by mean ±SEM. All other data represent mean ±SD.

### Supplemental Figure 6

A

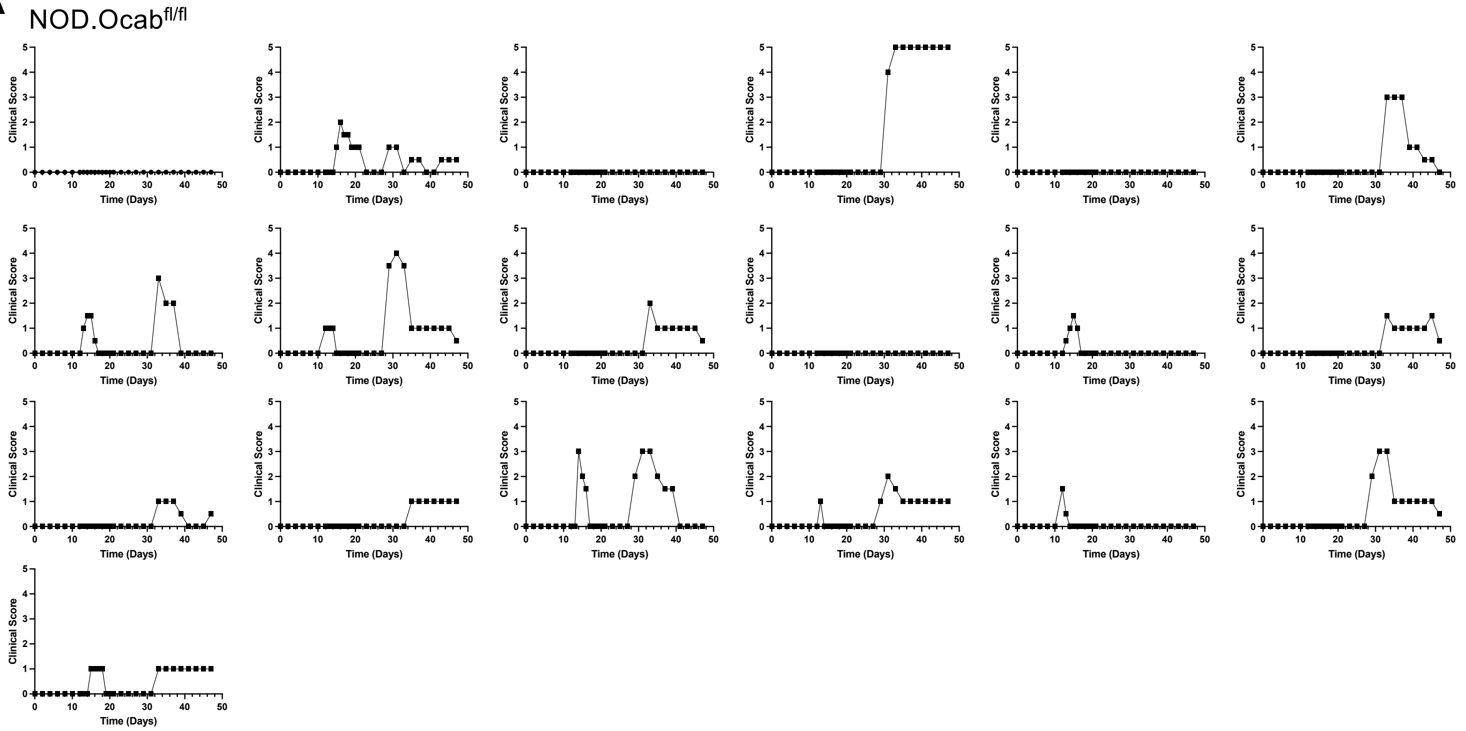


B

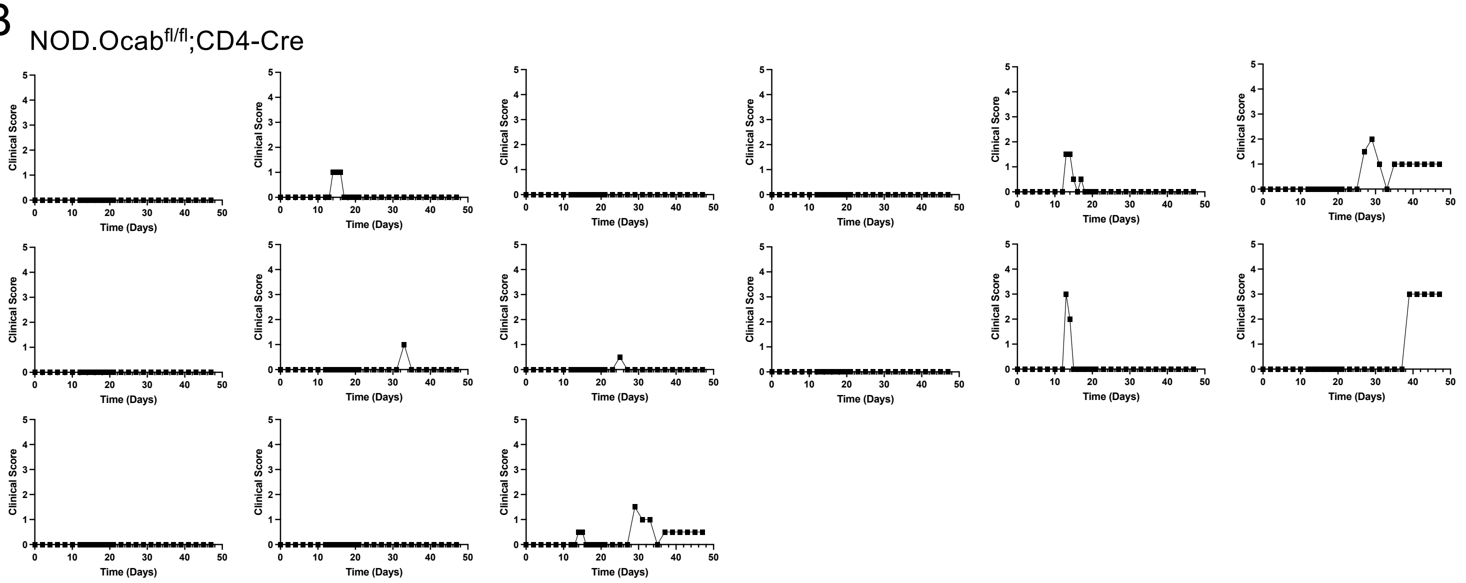

**Supplemental Figure 6. Individual NOD mouse EAE clinical scores.** (**A**) Clinical scores of individual NOD.Ocab$^{fl/fl}$ mice following EAE induction. (**B**) Clinical scores of individual NOD.Ocab$^{fl/fl}$;CD4-Cre mice following EAE induction.

Supplemental Figure 7

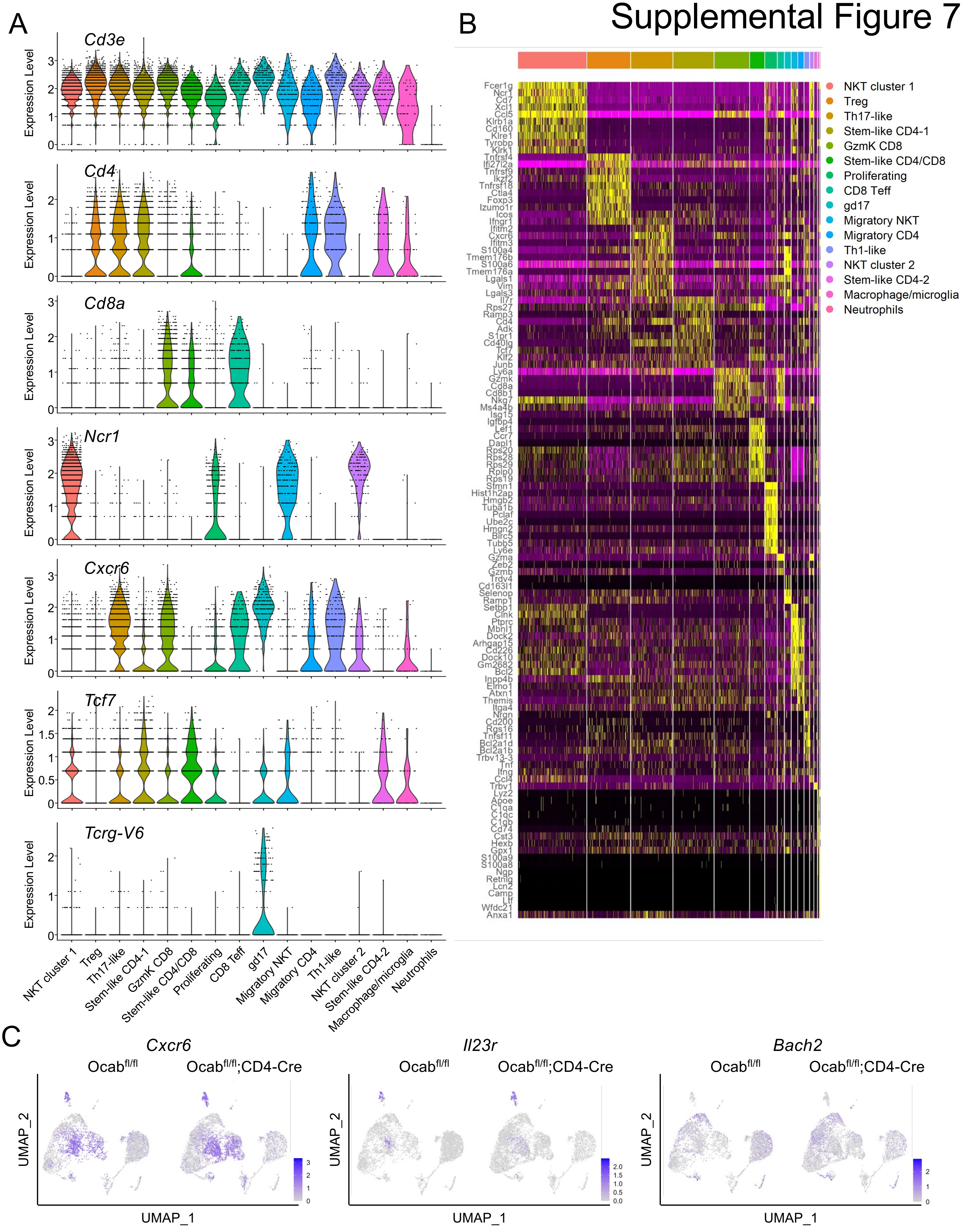

**Supplemental Figure 7. Additional remission scRNA-seq cluster annotation, heat map of gene enrichments by cluster, and additional feature plots.** (**A**) Violin plots showing cluster gene expression of *CD3e*, *Cd4*, *Cd8a*, *Ncr1*(NKp46), *Cxcr6*, *Tcf7*, and *Tcrg-V6* at EAE remission (**B**) Heatmap showing the top 10 genes expressed by cluster at remission EAE timepoint. (**C**) UMAP feature plots showing expression of *Cxcr6*, *Il23r*, and *Bach2* amongst clusters at remission.

### Supplemental Figure 8

A

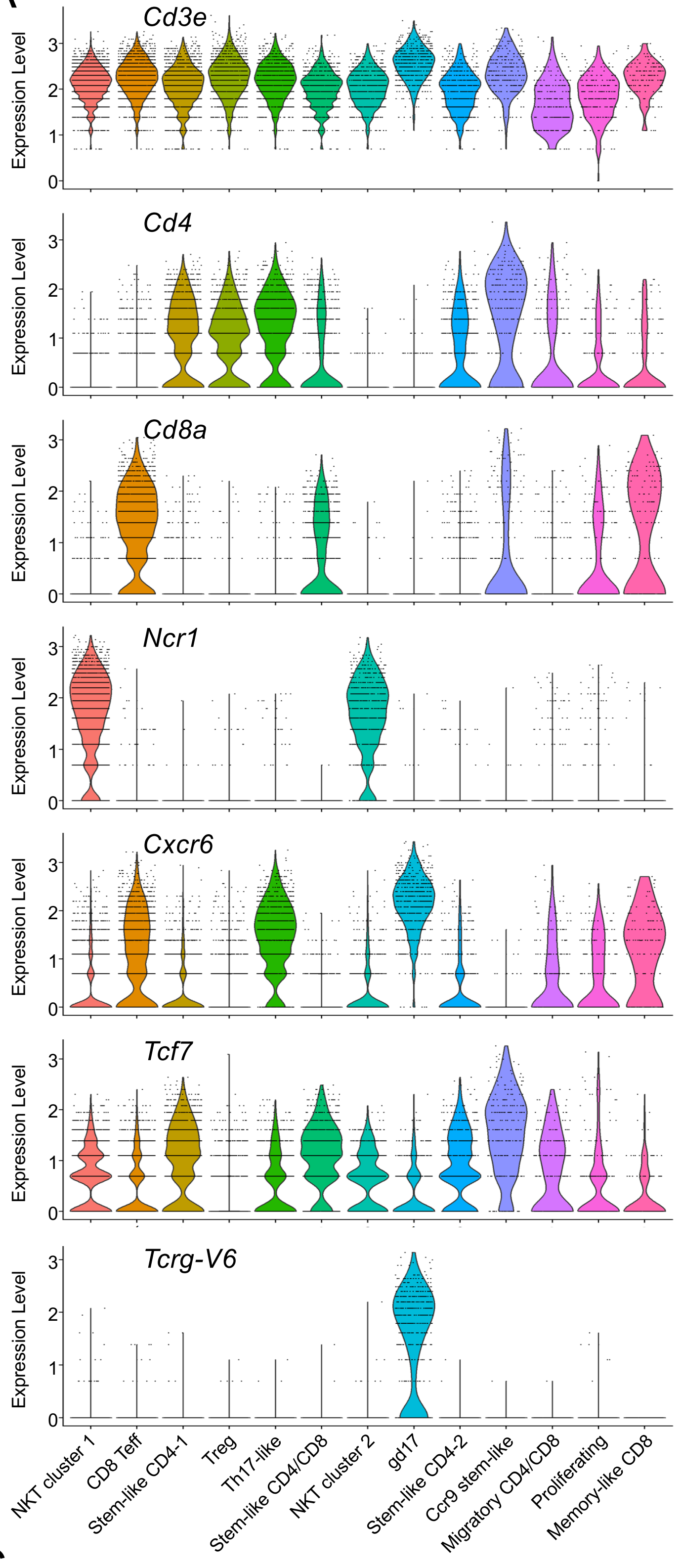


B

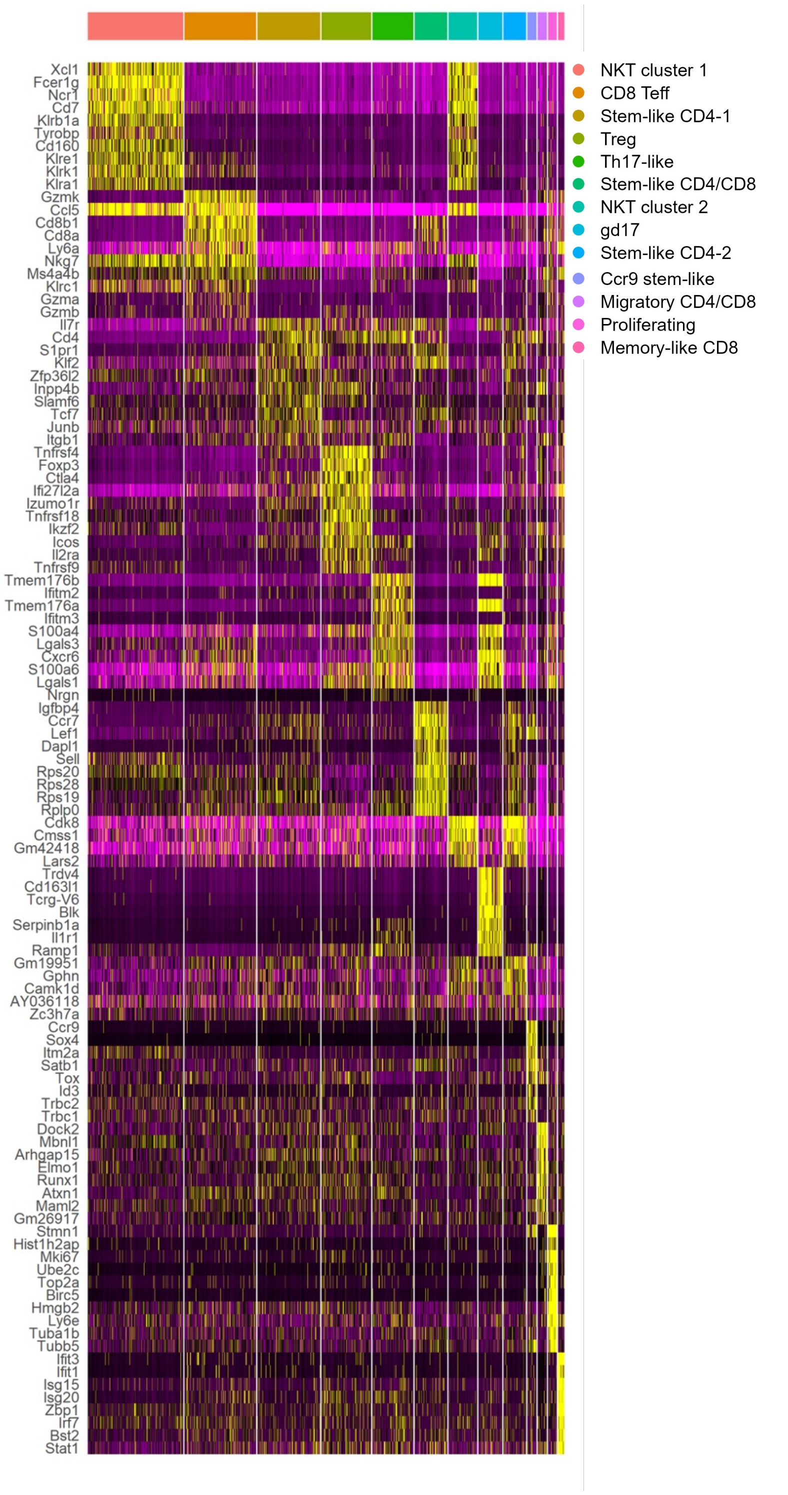


C

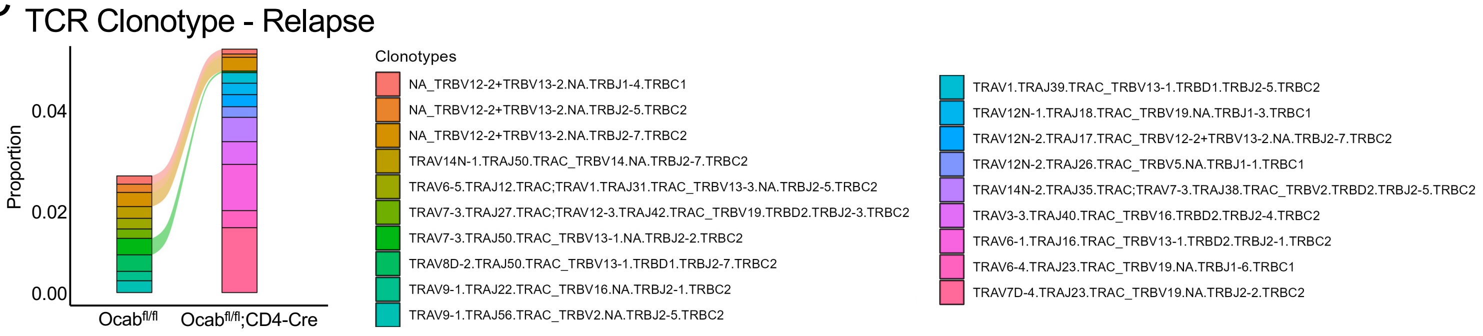

**Supplemental Figure 8. scRNA-seq cluster annotation genes, heat map of NOD.EAE gene enrichments by cluster, and TCR clonotype overlap between conditions at the relapse timepoint.** (**A**) Violin plots showing cluster gene expression of *CD3e*, *Cd4*, *Cd8a*, *Ncr1*(NKp46), *Cxcr6*, *Tcf7*, and *Tcrg-V6* at EAE relapse. (**B**) Heatmap showing the top 10 genes expressed by cluster at relapse EAE timepoint. (**C**) Alluvial plot showing top 10 TCR clonotype gene overlap between Ocab$^{fl/fl}$ and Ocab$^{fl/fl}$;CD4-Cre groups. (**C**) UMAP feature showing the expression of *CD4*, *CD8a, Tox*, *Sell*, *Ccr7*, and *Ccr4* amongst clusters at relapse timepoint.

### Supplemental Figure 9

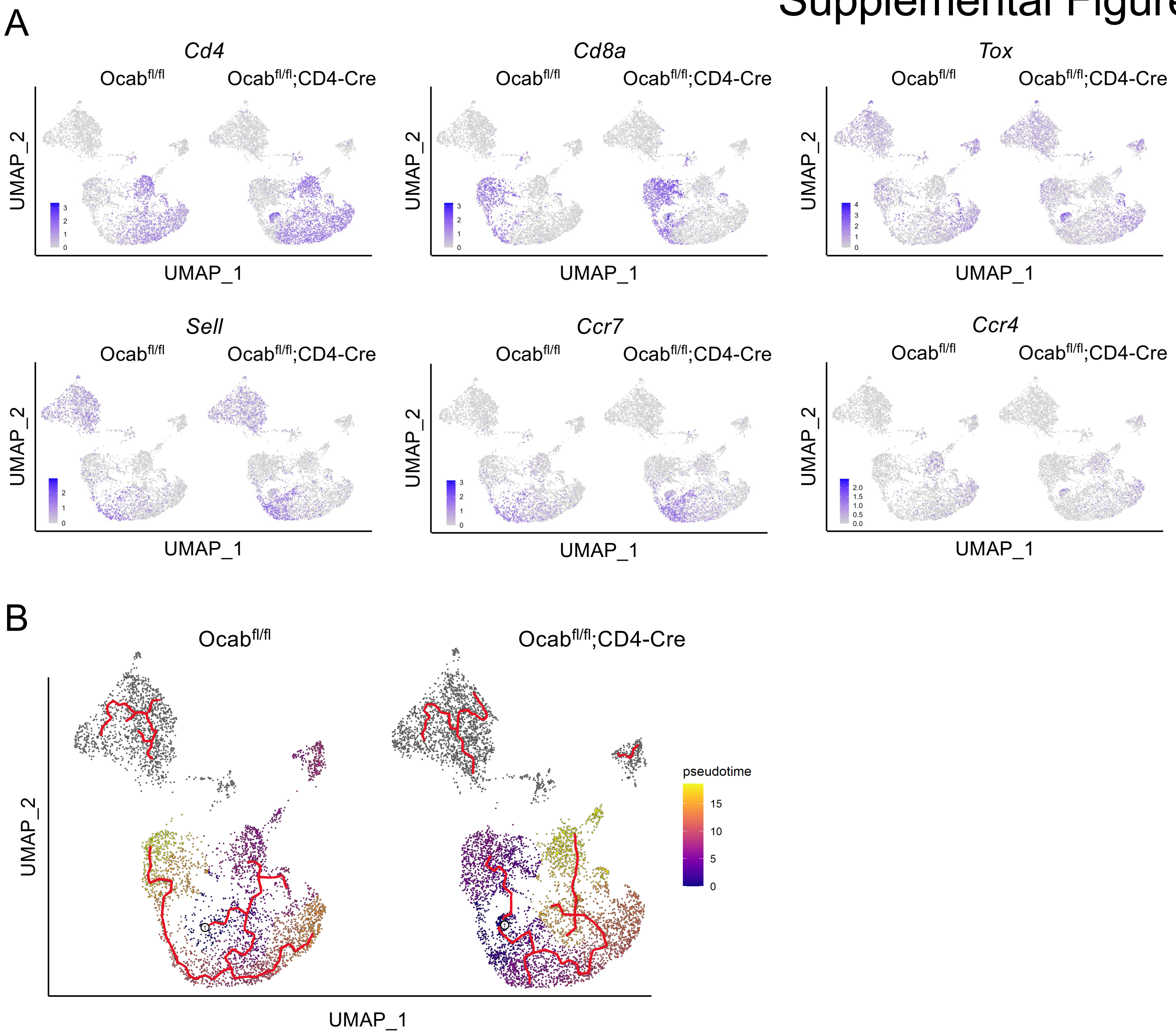

**Supplemental Figure 9. Example relapse feature plots and pseudotime analysis of relapse single cell RNA seq.** (**A**) UMAP feature showing the expression of *CD4*, *CD8a, Tox*, *Sell*, *Ccr7*, and *Ccr4* amongst clusters at relapse timepoint. (**B**) Pseudotime analysis of relapse single cell from root starting node at or near the *Ccr9*-expressing stem-like cluster.
